# An orally bioavailable 4-phenoxy-quinoline compound as a potent AURKB relocation blocker for cancer treatment

**DOI:** 10.1101/2023.01.29.526078

**Authors:** Jinhua Li, Ting Zhang, Qiong Shi, Gang Lv, Xiaohu Zhou, Namrta Choudhry, Julia Kalashova, Chenglu Yang, Hongmei Li, Yan Long, Balasubramaniyan Sakthivel, Naganna Nimishetti, Hong Liu, Thaddeus D. Allen, Jing Zhang, Dun Yang

## Abstract

We investigated a novel 4-phenoxy-quinoline-based scaffold that mislocalizes the essential mitotic kinase, AURKB. Here, we evaluated the impact of halogen substitutions (F, Cl, Br, I) on this scaffold with respect to various drug parameters. Br-substituted **LXY18** was found to be a potent and orally bioavailable disruptor of cell division, at sub-nanomolar concentrations. **LXY18** prevents cytokinesis by blocking AURKB relocalization in mitosis and exhibits broad-spectrum antimitotic activity *in vitro*. With a favorable PK profile, it shows widespread tissue distribution including the blood-brain barrier penetrance and effective accumulation in tumor tissues. More importantly, it markedly suppresses tumor growth. The novel mode of action of **LXY18** may eliminate some drawbacks of direct catalytic inhibition of AURKs. Successful development of **LXY18** as a clinical candidate for cancer treatment could enable a new, less toxic means of antimitotic attack that avoids drug resistance mechanisms.

---

The AURK family consists of three homologous serine/threonine kinases, AURKA, AURKB, and AURKC.^1^ They are key regulators of cell division and attractive oncology targets. Small molecule inhibitors of AURKs do not directly interfere with microtubules and a review of clinical studies suggests these inhibitors may avoid the peripheral neuro-toxicity observed with spindle toxins.^2^ Targeting these key mitotic kinases may, therefore, have advantages over standard chemotherapeutic approaches.

The AURKs play distinct roles in cell division. While AURKA is essential for the proper assembly of a bipolar mitotic spindle,^2^ AURKB and AURKC are catalytic subunits of the chromosomal passenger protein (CPP) complex, which coordinates karyokinesis with the completion of cytokinesis.^3^ AURKC, however, is a meiotic, rather than mitotic kinase, so AURKB is the catalytic subunit in somatic cells. The localization of the CPP complex is highly coordinated during mitosis. For example, the CPP complex is dynamically relocalized from chromosomes to the spindle midzone with the onset of anaphase. Preventing relocalization may just as effectively compromise AURKB function as catalytic inhibition, but with the added bonus of not altering kinase function during interphase or in non-dividing cells. Despite this, no therapeutic is currently known to block the mitotic localization of AURKB for cancer treatment. Instead, various catalytic inhibitors of AURKs have been clinically tested. The AURK inhibitors in clinical trials often inhibit all three AURKs without much discrimination,^4^ and several trials have been terminated partly due to adverse effects that stemmed from off-target interactions with other kinase families. Also, drug resistance due to compensating AURK mutation has been suggested to contribute to a lack of durable efficacy in clinical trials.^5^

We previously described a mechanism-informed, phenotypic screening (MIPS) assay of a synthetic compound library,^6,7^ for compounds that disrupt the localization, but not catalytic activity, of the CPP complex. Such compounds prevent cytokinesis, induce polyploidy, and trigger cell death in model rodent and human-derived cancer cell lines. We incorporated phenotypic screening into an optimization protocol and derived a scaffold where compounds shared the ability to disrupt CPP complex localization to varying degrees. These 4-phenoxy-quinoline derivatives exhibited anticancer activity in a variety of human cancer cell lines *in vitro*.^7^

We chose one compound, **LG182**, which demonstrated potent *in vitro* activity and limited *in vivo* activity,^7^ for further optimization, with the goal of finding a compound with adequate pharmacokinetic (PK) and pharmacodynamic (PD) properties for development as a new drug candidate. This starting compound was subjected to iterative rounds of structural modification coupled with phenotypic activity screening. Halogen analogs of **LG182** were found to improve the compound. Halogenation is often used in late-stage lead optimization, improving drug metrics,^8^ and sometimes even increasing binding affinity through the formation of an additional halogen bond.^9^ Here we present studies on a panel of compounds with halogen substitutions in place of an OMe functional group. Bromine substitution produced a promising compound, which we characterize here as **LXY18**.

Previous work with these 4-phenoxy-quinoline-based derivatives focused on structure-activity relationships that could be discerned from systemically modifying side groups attached to varying locations of the 3-ring core **(Scheme 1)**.^6,7^ Our potent *in vitro* compound, **LG182**, harbors an aromatic ring with di-meta substitutions of an OMe group and an acetamide group. Leaving this structural feature alone, we explored substitutions at the position labeled R of the ring most distal to the di-meta ring. In **LG182**, this site is occupied by an OMe group. However, many other substitutions at R were not well tolerated. Since the presence or absence of the Ome group did impact potency, we surmised substitution at this site was important but maybe size limited. In addition, positive contribution of fluorine to the potency prompted us to explore the other halogen members at this position. These additional analogs were synthesized as described (**Scheme 1**).

**Scheme 1.**
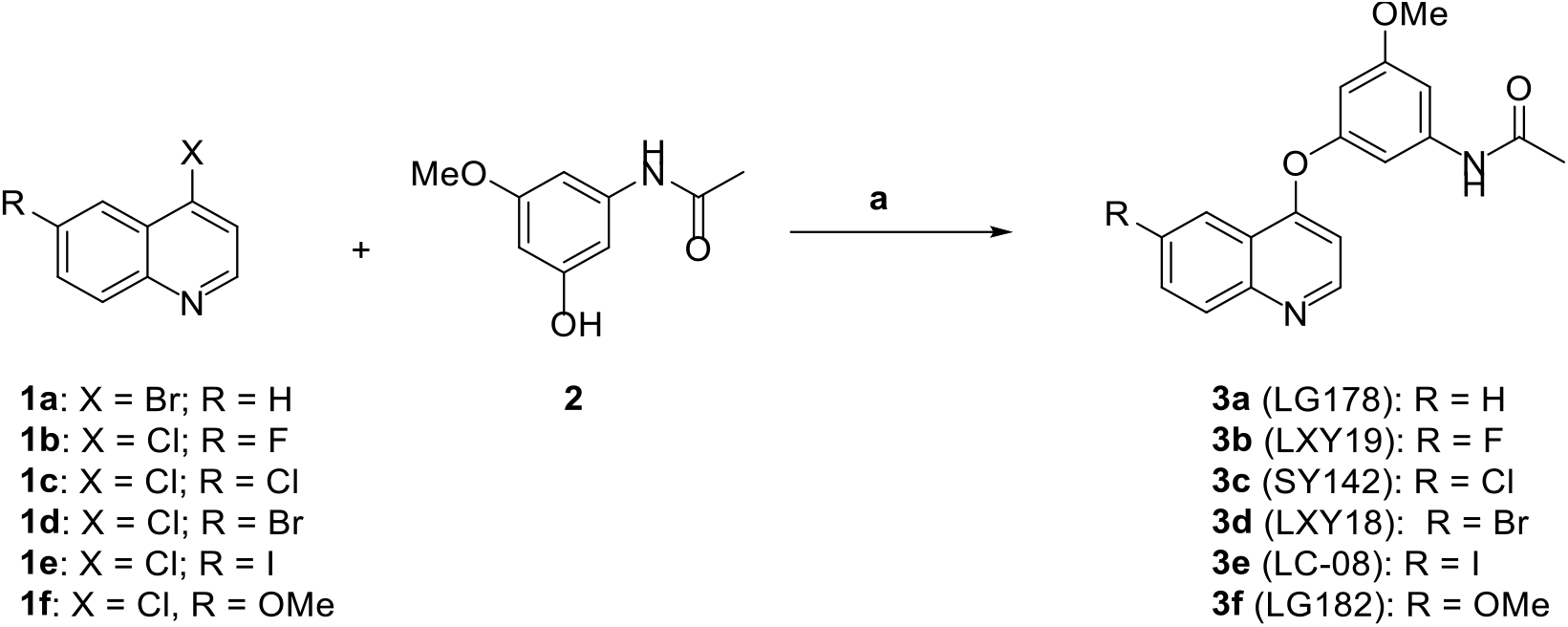
Synthesis of 6-Halo, 4-Phenoxy Quinoline Derivatives^*a a*^Reagents and conditions are as follows: (a) K_2_CO_3_, DMF, N_2_, 115 °C, 12 h, 50-89%.

The impact of halogen substitutions at R on the drug-like properties of the molecules was first computationally modeled. All of these halogen-substituted compounds satisfied the rule of five,^10^ which predicts the oral availability of drugs (**Table S1**). Consistent with the notion that halogen substituents make molecules more lipophilic, the substitution of hydrogen at R1 with F, Cl, Br, or I, gradually increased cLog P, a calculated measure of compound hydrophilicity, from 3.99 to 4.13, 4.65, 4.76 and 5.01 respectively. Four compounds with different halogen substitutions and the parental compound exhibited the same topological polar surface area (TPSA) value of 60.45 (**Table S1**). They are all predicted to be able to cross the blood-brain barrier since a compound with a TPSA value less than 90 Å^2^ correlates with an increased ability to transition across the blood-brain barrier.^11^ All of these compounds were stable in mouse plasma and human serum (**Table S2**) or in a buffer with a pH value of 2.2 or 7.4 (**Table S3**). Halogen substitutions did not affect solubility much since all analogs had similar solubility. Consistent with the presence of a quinoline ring, solubility was pH-dependent with >10-fold higher kinetic solubility at pH 2.2 relative to pH 7.4 for each compound (**Table 3**). In accordance with the pH-dependent solubility, the experimentally determined Log D for each compound decreased >10-fold when pH declined from 7.4 to 2.2. We next determined the bioactivity of these quinoline analogs.

Retinal pigment epithelial (RPE) cells expressing the MYC oncoprotein, plus the histone 2B protein fused to a green fluorescent protein (H2B-GFP) were used to assess drug potency *in vitro*. The minimum effective concentration (MEC) in RPE-MYC^H2B-GFP^ cells that induces phenotypes reminiscent of AURKB inhibition was used to compare compounds, as this was previously found to be an effective metric.^6,7^ The most easily discernable outcome from inhibiting the localization of the chromosomal passenger protein complex is the induction of polyploid cells, as AURKB localizes during telophase to the midbody and mediates the induction of cytokinesis.^12^ However, the MEC for mitotic arrest, polyploidy, and cell death were the same in this assay as we observed before,^7^ and we referred to them as MEC without discrimination (**Table 1)**. Substitution with either of three heavier halogens, Cl, Br, and I, increased the potency to a similar extent, reducing MEC 3 to 4-fold, relative to OMe-substituted **LG182** and F-substituted **LXY19**.

**Table 1.**
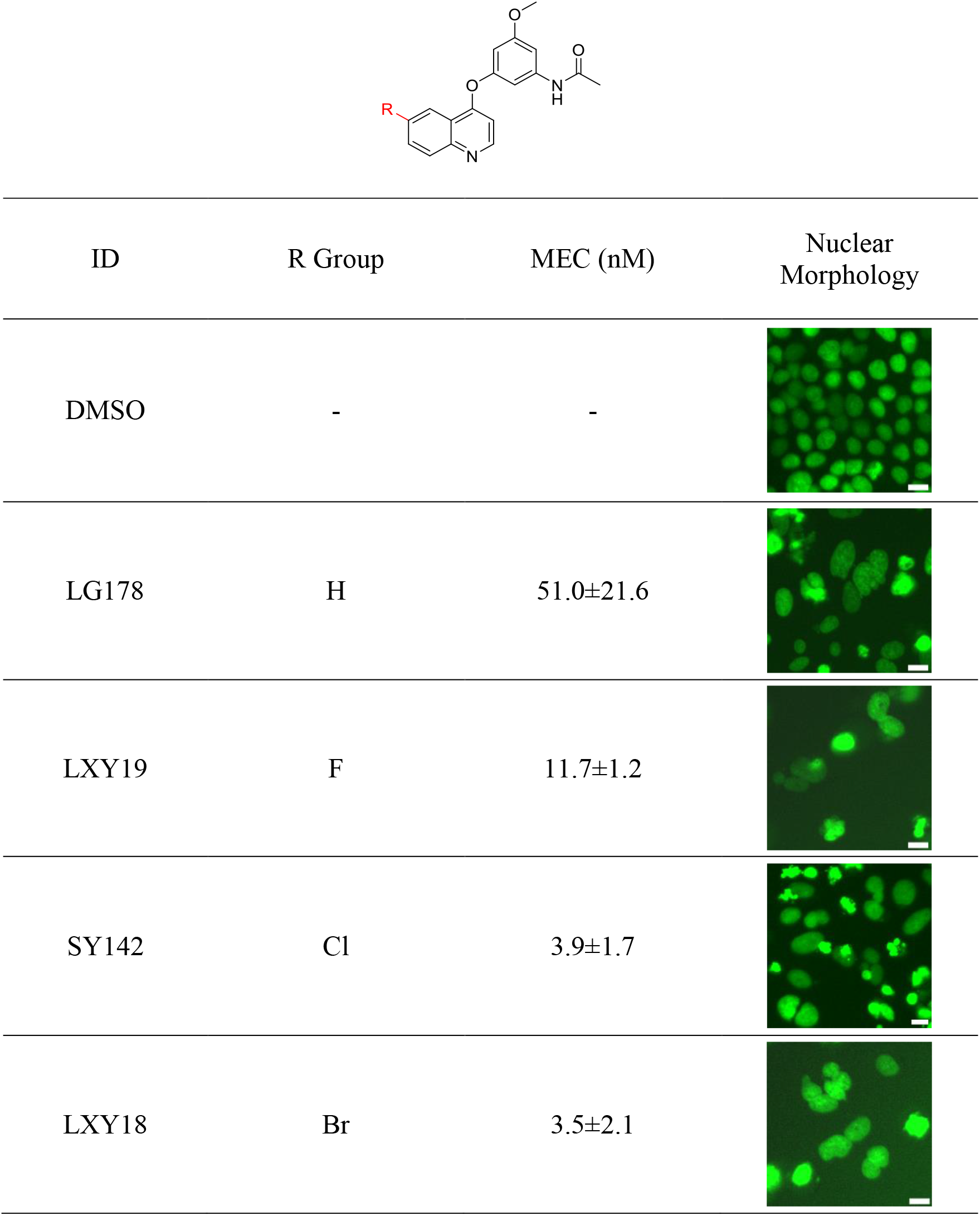
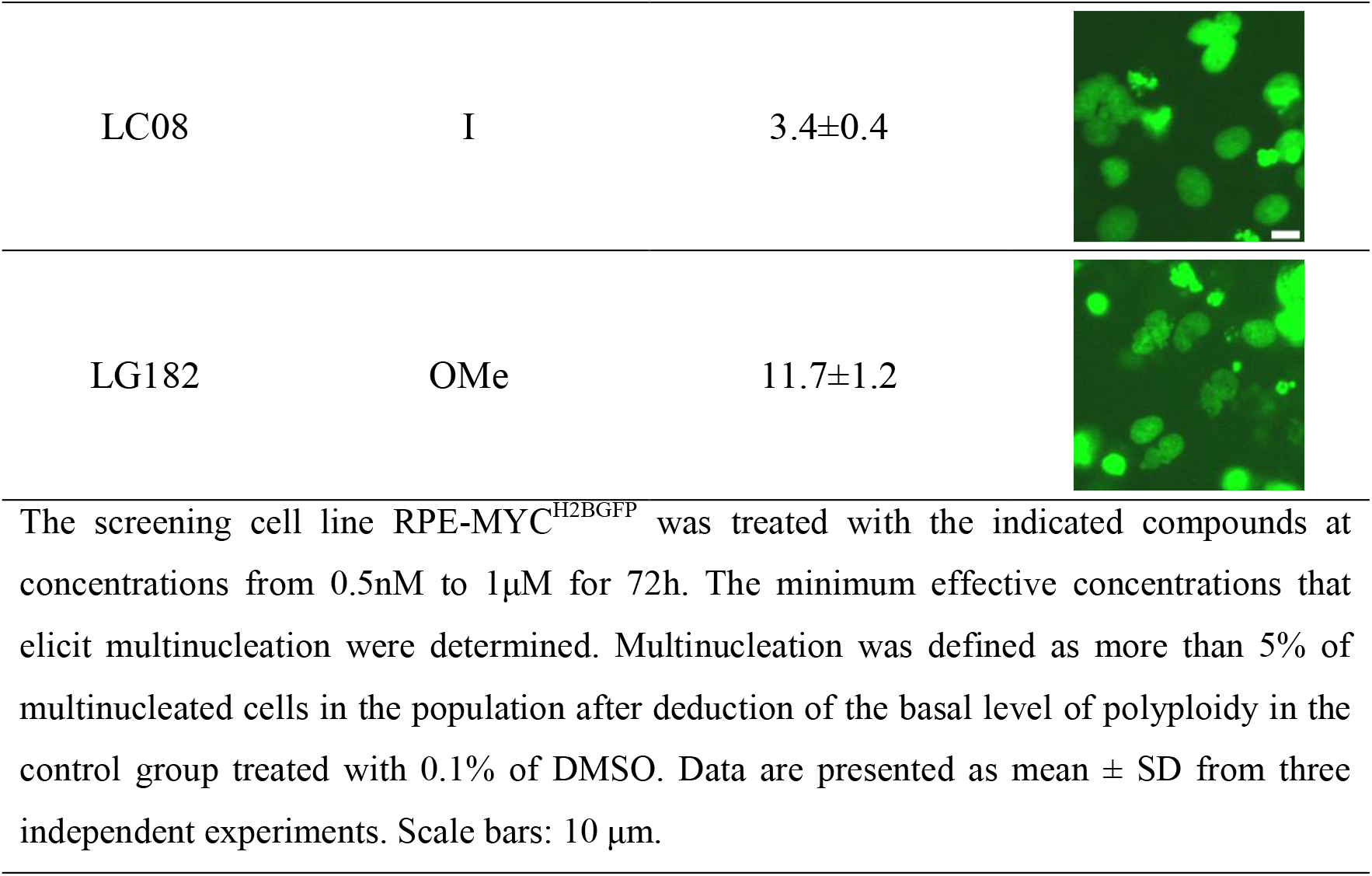
The minimum effective concentrations (MEC) of quinoline analogs to induce polyploidy.

| ID | R Group | MEC (nM) | Nuclear Morphology |
| --- | --- | --- | --- |
| DMSO | - | - |  |
| LG178 | H | 51.0±21.6 |  |
| LXY19 | F | 11.7±1.2 |  |
| SY142 | Cl | 3.9±1.7 |  |
| LXY18 | Br | 3.5±2.1 |  |
| LC08 | I | 3.4±0.4 |  |
| LG182 | OMe | 11.7±1.2 |  |

The screening cell line RPE-MYC<sup>H2BGFP</sup> was treated with the indicated compounds at concentrations from 0.5nM to 1μM for 72h. The minimum effective concentrations that elicit multinucleation were determined. Multinucleation was defined as more than 5% of multinucleated cells in the population after deduction of the basal level of polyploidy in the control group treated with 0.1% of DMSO. Data are presented as mean ± SD from three independent experiments. Scale bars: 10 μm.

Immunofluorescent (IF) analysis with markers that are used as surrogates for the activity of AURKs can be used to assess the catalytic inhibition of these mitotic kinases as we described before.^6,7^ For AURKB, the intensity of Histone 3 phosphorylation at Serine 10 (H3Ser10P), a direct target of AURKB in mitosis, is a well-established surrogate of its kinase activity. We have previously combined phenotypic screening in RPE-MYC^H2B-GFP^ cells with IF for surrogate markers of AURK activity as a MIPS method.^6,7^ Here, we continued to assess compounds with this combination approach to ensure the halogenated compounds are not catalytic inhibitors but added an IF assay to ensure they also disrupt AURKB positioning at the spindle midzone during anaphase. Halogenated compounds did so, irrespective of which halogen was substituted at R. IF examination of earlier stages of mitosis suggested that localization of AURKB on prometaphase kinetochores was not affected in two human cancer cell lines NCI-H23 and DU-145 and a model cell line RPEMYC^Bcl2^ (**Figure S1)**. Furthermore, AURKB was still present on anaphase chromosomes in all three cell lines (**Figures 1A** and **S2**), suggesting the compounds prevented the release of AURKB from chromosomes at the anaphase onset. Consequently, these compounds prevented repositioning at the spindle midzone, so despite our previous designation of this class of compounds as AURKB localization inhibitors, these compounds may be better described as inhibitors of AURKB relocation, as earlier localization during mitosis does not seem to be altered. The lowest concentrations of these compounds that prevented relocation were equivalent to MEC in the RPE-MYC^H2B-GFP^ assay, so we concluded that cell-division defects elicited by these compounds could be attributed to a failure of an essential AURKB function. None of these compounds inhibited phosphorylation of the AURKB substrate Histone 3 at Serine 10 (H3Ser10P) (**Figures 1B** and **S3, Table S4**) or AURKA autophosphorylation at Thr288 (**Table S4** and **Figures S3** and **S4**), so these compounds do not inhibit the catalytic activity of AURKs.

**Figure 1.**
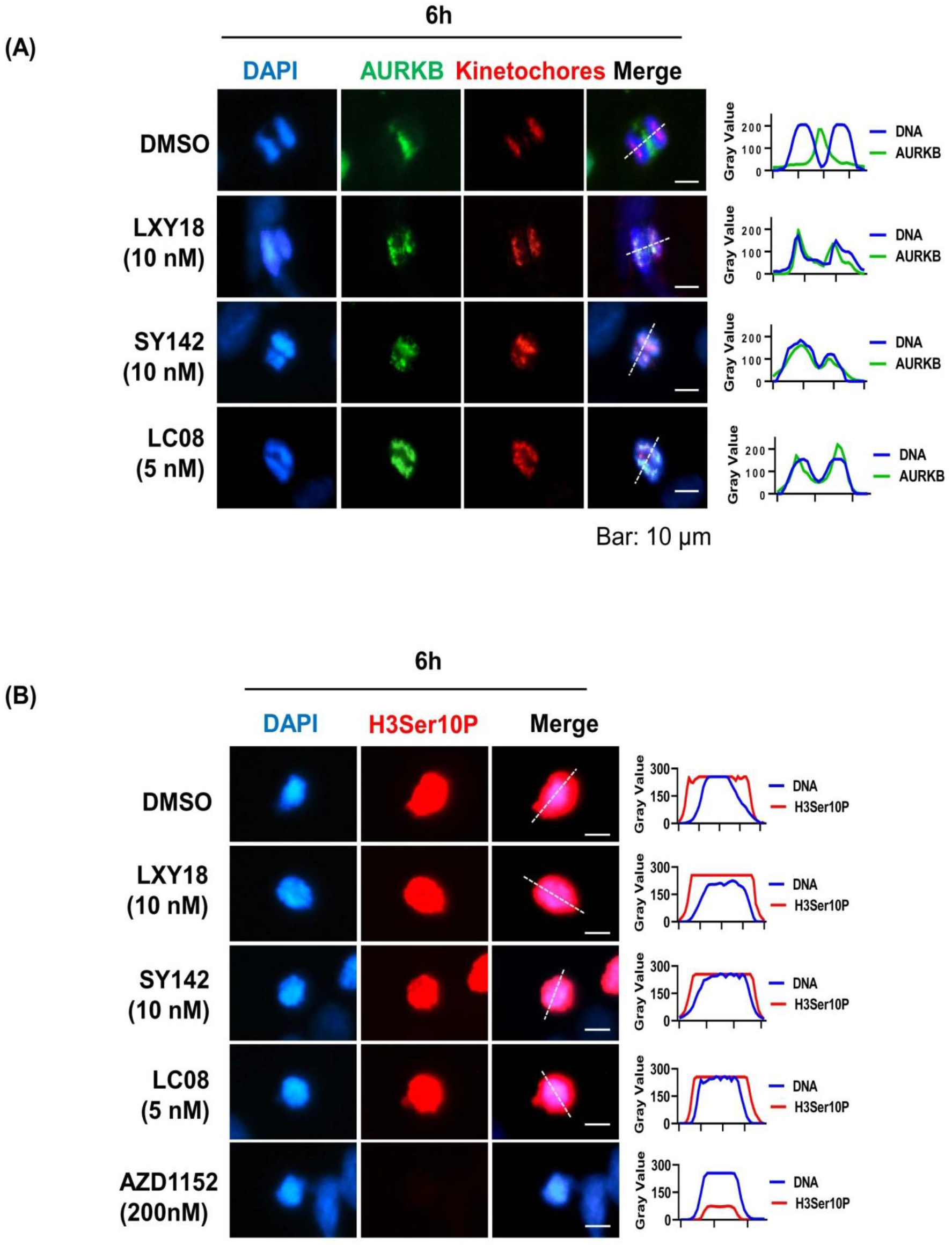
The effect of quinoline analogs on kinase activity and localization of AURKB. RPE-MYC^BCL2^ cells were treated for 6 h with the indicated compounds before stained with DAPI alongside antibodies that recognize H3Ser10P, AURKB, or kinetochore proteins. Cells treated with vehicle (0.1% of DMSO) were used as a negative control. (A) The effect on AURKB mitotic localization. (B) The effect on phosphorylation of H3 at Ser10. For quantitation in A and B, the fluorescence intensity along the white dashed line was determined using ImageJ software and was plotted as a relative gray value over distance in pixels. More than 20 mitotic cells in each group were scored and all displayed similar images as the representative cell presented. Scale bars: 10 μm.

To confirm these halogenated compounds have a broad anticancer activity, we assayed their bioactivities in a panel of 17 cancer cell lines from various tissue origins. The formation of multinucleation was used as a convenient readout for the disablement of AURKB.^7^ Consistent with the findings in the screening assay, **SY142, LXY18**, and **LC08** had lower MEC in inducing polyploidy relative to **LG182** and **LXY19** in the majority of the human cancer cell lines (**Figure 2**), suggesting that substitutions with heavier halogens improved potency in general in human cancer cell lines.

**Figure 2.**
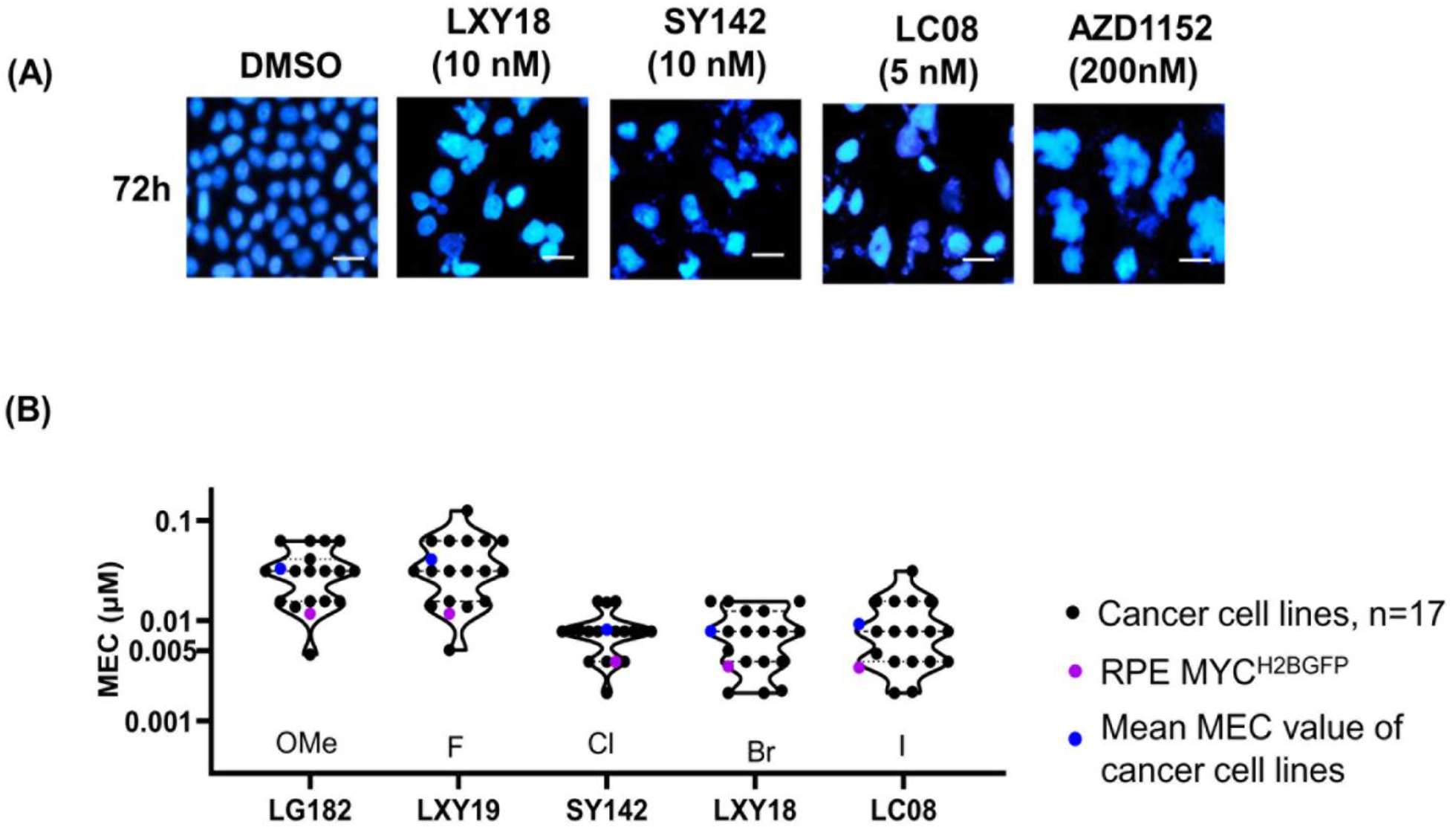
The induction of multinucleation by quinoline analogs. RPE-MYC^BCL2^ cells and 17 human cancer cell lines were treated for 72 h with the indicated compounds at the indicated concentrations (A) or a concentration range from 0.5 nM to 1 μM (B). Cells were then stained with DAPI to visually assess the minimum effective concentrations (MEC) using multinucleation as a metric of activity. A. Induction of multinucleation. scale bars: 10 μm. B.Violin plot of the minimum effective concentrations for polyploidy. Each black dot denotes a cancer cell line and the RPE-MYC^H2B-GFP^ cell line is in purple. The blue dots indicate the mean of MEC for a given compound in 17 cancer cell lines.

Halogenation of sp^2^ carbons is known to increase lipophilicity and improve membrane permeability and oral absorption.^13^ We next examined whether halogenation impacted plasma drug exposure. Compounds were delivered to mice orally using three types of delivery vehicles. We tested lipid-soluble corn oil, hydrophilic hydroxypropyl methylcellulose (HPMC), and water-soluble, neutral polyethylene glycol 300 (PEG300), which represent oil, suspension, and aqueous delivery modes, respectively. Plasma samples were collected from 0 to 6 hours and drug concentration was quantified by LC-MS/MS (schematic **Figure 3A)**. Among the compounds tested, **LXY18** achieved the highest level of plasma exposure with each of the three formulations, as judged by dose-normalized AUC_0-6h_ (**Figure 3B**) or the AUC_0-6h_ after further normalization to bioactivity (**Figure 3C**). Among the three oral delivery formulations, the largest AUC or C_max_ value for **LXY18** was obtained with lipid-soluble corn oil (**Figure 3B-E**).

**Figure 3.**
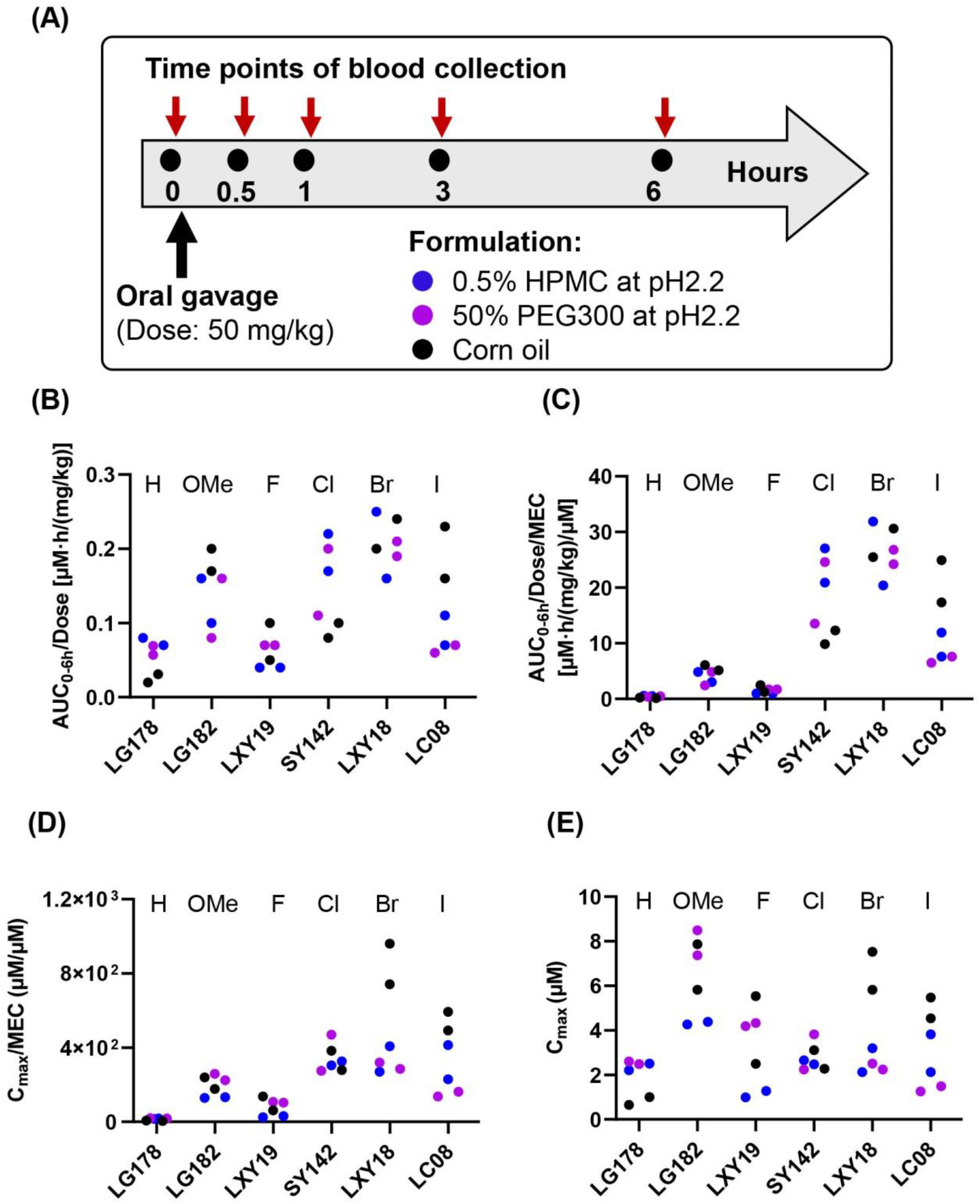
Pharmacokinetic parameters of quinoline analogs after oral gavage. Compounds were suspended or dissolved into the indicated formulation and delivered to nude mice (8-11 weeks old female BAL B/c). Two mice were used for each formulation. (A) Schematic diagram of timeline for blood collection after a single oral administration. (B) Scatter plot of the dose-normalized AUC_0-6h_ value. The area under the plasma concentration-time curve from time zero to 6 h was divided by the dose. (C) The scatter plot of the AUC_0-6h_/Dose/MEC (Dose-normalized AUC_0-6h_ divided by the mean of MEC). (D) The scatter plot of the activity normalized C_max_ (C_max_/MEC). (E) The scatter plot of the maximum plasma concentration (C_max_) after a single dose of 50 mg/kg. Each dot in B-E represents a mouse. The color indicates the formulation described in A.

Above findings suggested that **LXY18** had both improved potency and plasma exposure after an oral dose and was a top compound among the halogenated analogs. The pharmacokinetic parameters of **LXY18** were studied further in Wistar rats. Data are summarized in **Table 2**. After oral administration of 2 mg/kg of **LXY18** in corn oil, we observed a peak absorption (C_max_) of 0.46 ± 0.35 μM 1 h post-administration, a half-life (T_1/2_) of 2.84 ± 1.36 hour. Compared with the AUC of the same amount of **LXY18** intravenously delivered, the overall oral availability was calculated as 51.40%.

**Table 2.**
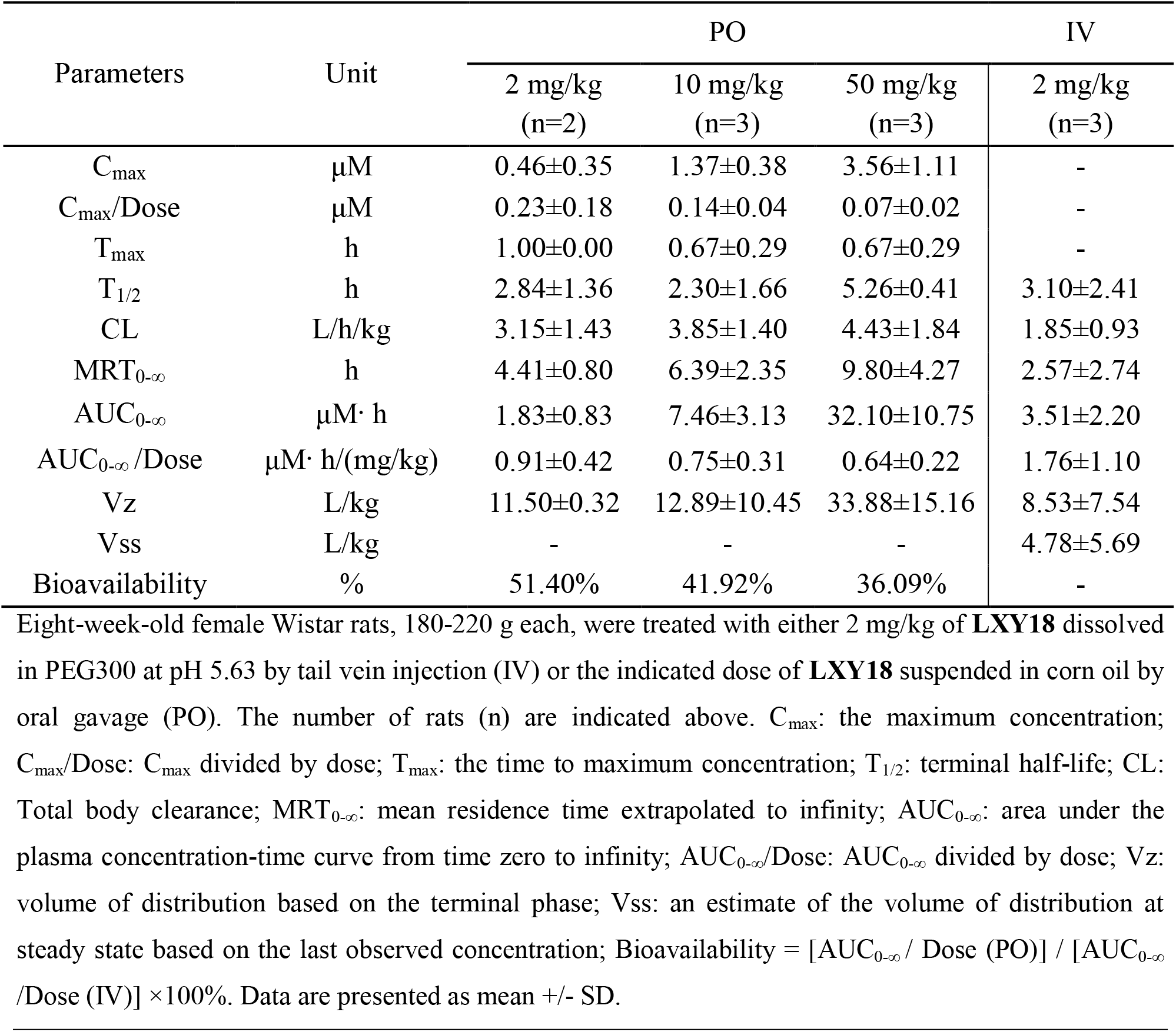
Pharmacokinetic parameters of LXY18 after a single oral or intravenous administration in rats.

| Parameters | Unit | PO |  |  | IV |
| --- | --- | --- | --- | --- | --- |
|  |  | 2 mg/kg<br>(n=2) | 10 mg/kg<br>(n=3) | 50 mg/kg<br>(n=3) | 2 mg/kg<br>(n=3) |
| C<sub>max</sub> | μM | 0.46±0.35 | 1.37±0.38 | 3.56±1.11 | - |
| C<sub>max</sub>/Dose | μM | 0.23±0.18 | 0.14±0.04 | 0.07±0.02 | - |
| T<sub>max</sub> | h | 1.00±0.00 | 0.67±0.29 | 0.67±0.29 | - |
| T<sub>1/2</sub> | h | 2.84±1.36 | 2.30±1.66 | 5.26±0.41 | 3.10±2.41 |
| CL | L/h/kg | 3.15±1.43 | 3.85±1.40 | 4.43±1.84 | 1.85±0.93 |
| MRT<sub>0-∞</sub> | h | 4.41±0.80 | 6.39±2.35 | 9.80±4.27 | 2.57±2.74 |
| AUC<sub>0-∞</sub> | μM·h | 1.83±0.83 | 7.46±3.13 | 32.10±10.75 | 3.51±2.20 |
| AUC<sub>0-∞</sub>/Dose | μM·h/(mg/kg) | 0.91±0.42 | 0.75±0.31 | 0.64±0.22 | 1.76±1.10 |
| Vz | L/kg | 11.50±0.32 | 12.89±10.45 | 33.88±15.16 | 8.53±7.54 |
| Vss | L/kg | - | - | - | 4.78±5.69 |
| Bioavailability | % | 51.40% | 41.92% | 36.09% | - |

Eight-week-old female Wistar rats, 180-220 g each, were treated with either 2 mg/kg of **LXY18** dissolved in PEG300 at pH 5.63 by tail vein injection (IV) or the indicated dose of **LXY18** suspended in corn oil by oral gavage (PO). The number of rats (n) are indicated above. C<sub>max</sub>: the maximum concentration; C<sub>max</sub>/Dose: C<sub>max</sub> divided by dose; T<sub>max</sub>: the time to maximum concentration; T<sub>1/2</sub>: terminal half-life; CL: Total body clearance; MRT<sub>0-∞</sub>: mean residence time extrapolated to infinity; AUC<sub>0-∞</sub>: area under the plasma concentration-time curve from time zero to infinity; AUC<sub>0-∞</sub>/Dose: AUC<sub>0-∞</sub> divided by dose; Vz: volume of distribution based on the terminal phase; Vss: an estimate of the volume of distribution at steady state based on the last observed concentration; Bioavailability = [AUC<sub>0-∞</sub> / Dose (PO)] / [AUC<sub>0-∞</sub> / Dose (IV)] × 100%. Data are presented as mean +/- SD.

Next, three oral doses 2, 10, and 50 mg/kg of **LXY18** were compared. Dose-linear increases in AUC and Cmax were observed. The bioavailability decreased from 51.40% to 41.92% and 36.09% when the dose of **LXY18** was elevated from 2 to 10 and 50 mg/kg. Rats that received two lower oral doses of **LXY18** had a similar apparent volume of distribution during the terminal phase (Vz) of 11.50-12.89 L/kg and a half-life of 2.30-2.84 h. The half-life increased to 5.26 h and Vz was elevated to 33.88 L/kg in rats that were treated with 50 mg/kg. After the tail vein injection of **LXY18**, the terminal phase Vz and steady-state volume of distribution (Vss) were determined as 8.53 ± 7.54 L/kg and 4.78 ± 5.69 L/kg, respectively (**Table 2**). The compound was widely distributed since the Vz or Vss value was more than six-fold larger than the rat’s total body water volume (0.7 L/kg).

Elucidation of drug tissue distribution is usually exploited to confirm suitable exposure for inhibition of the therapeutic target and can also identify potential tissues where toxicity may be problematic. Quantitative tissue distribution studies (**Figure 4**) were pursued to confirm the wide drug tissue distribution predicted by Vz and Vss values (**Table 2**). Tissues were dissected at 1, 4, or 8 h after a single oral administration of **LXY18** (**Figure 4A**). A plasma **LXY18** concentration of 3.22 ± 0.97 μM was found 1 h after oral administration and **LXY18** was detected in all tissues and organs examined (**Figure 4B**). The highest **LXY18** exposure was present in adipose tissue (AUC_1-8h_ = 8.28 ± 2.19 μM·h). The liver had the lowest **LXY18** exposure (AUC_1-8h_ = 0.87 ± 0.54 μM· h), likely resulting from metabolic clearance of the compound in the liver. Importantly, **LXY18** was detected in brain tissue and the CSF after 1 h with a concentration of 0.59 ± 0.27 μM and 0.02 ± 0.00 μM, respectively, suggesting that the compound transits the blood-brain barrier readily.

**Figure 4.**
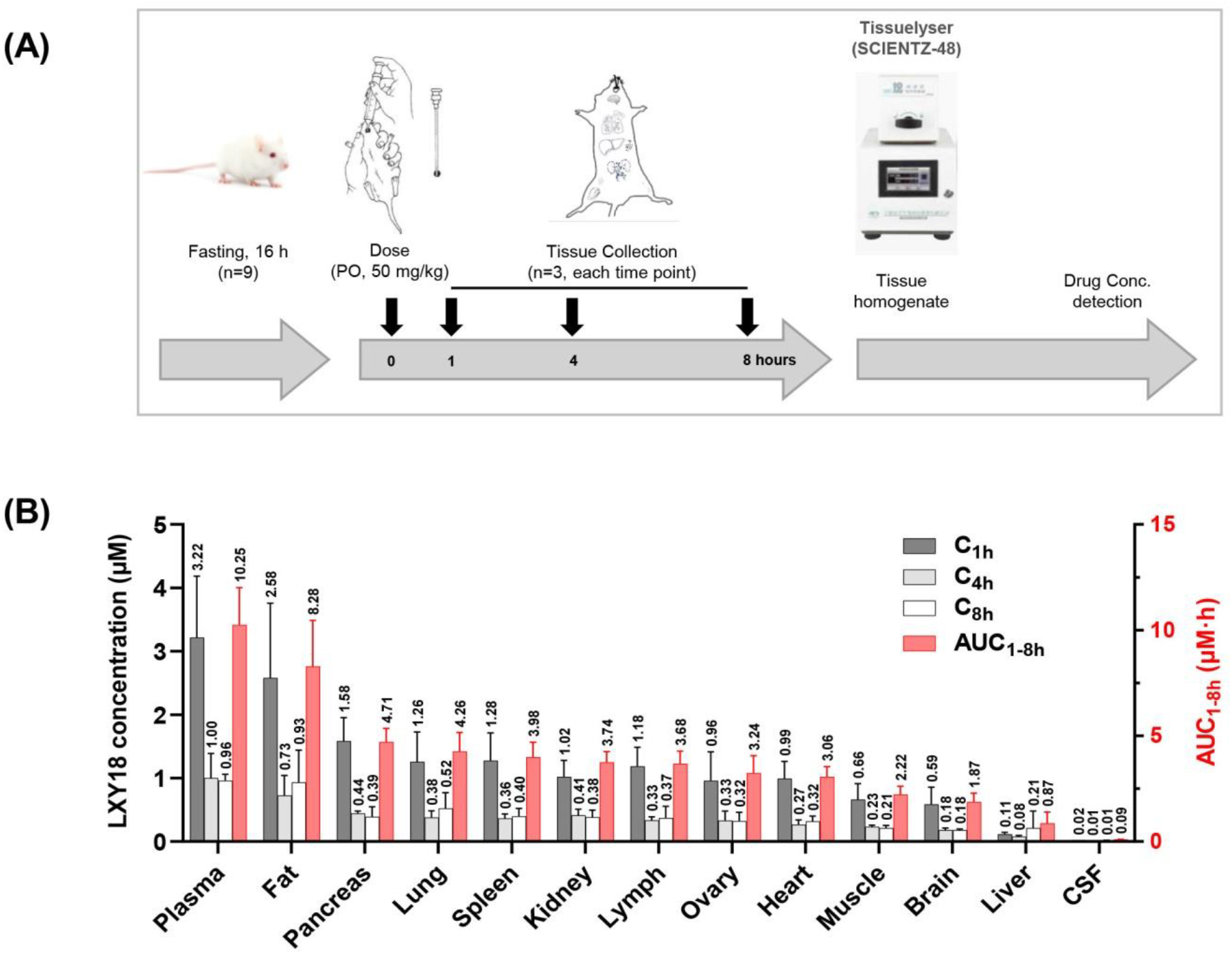
**LXY18** tissue distribution studies. Wistar rats received a single oral dose of 50 mg/kg **LXY18**. The peripheral blood was collected at 1, 4, and 8 h after drug administration. The plasma drug concentration at each time point was determined by LC-MS/MS. Three 9-week-old female Wistar rats, 200-230 g each, were used at each time point. (A) Flow chart of the tissue collection scheme for drug distribution studies. (B) Drug concentrations in different tissues. Besides the plasma drug concentrations at 1, 4, and 8 h (left y-axis), the area under the concentration-time curve from time 1 to 8 h (AUC_1-8h_) is also presented (right y-axis).

Plasma concentration has generally been considered a surrogate for the concentration of drug available to reach a tumor. However, intra-tumoral drug concentrations can vary considerably from plasma levels due to heterogeneity in the tumor microenvironment.^14^ Intratumoral concentration of **LXY18** was further tested in immunodeficient mice bearing xenografts of two different human cancer cell lines. In this experiment, we quantified **LXY18** after mice had received repeated treatments (**Figure 5A**). This design is to examine if **LXY18** might induce metabolic enzymes and consequently reduce **LXY18**’s plasma exposure after multiple treatments. The maximum tolerated dose (MTD) of **LXY18** orally delivered in corn oil was determined to be 100 mg/kg/day in BALB/c nude mice under a long-term daily repeat treatment. We chose to perform treatment with 50 mg/kg of **LXY18** twice a day for five days before **LXY18** was quantified in the plasma after the last oral treatment. Tumor tissues were also harvested 6 hours after the last treatment.

**Figure 5.**
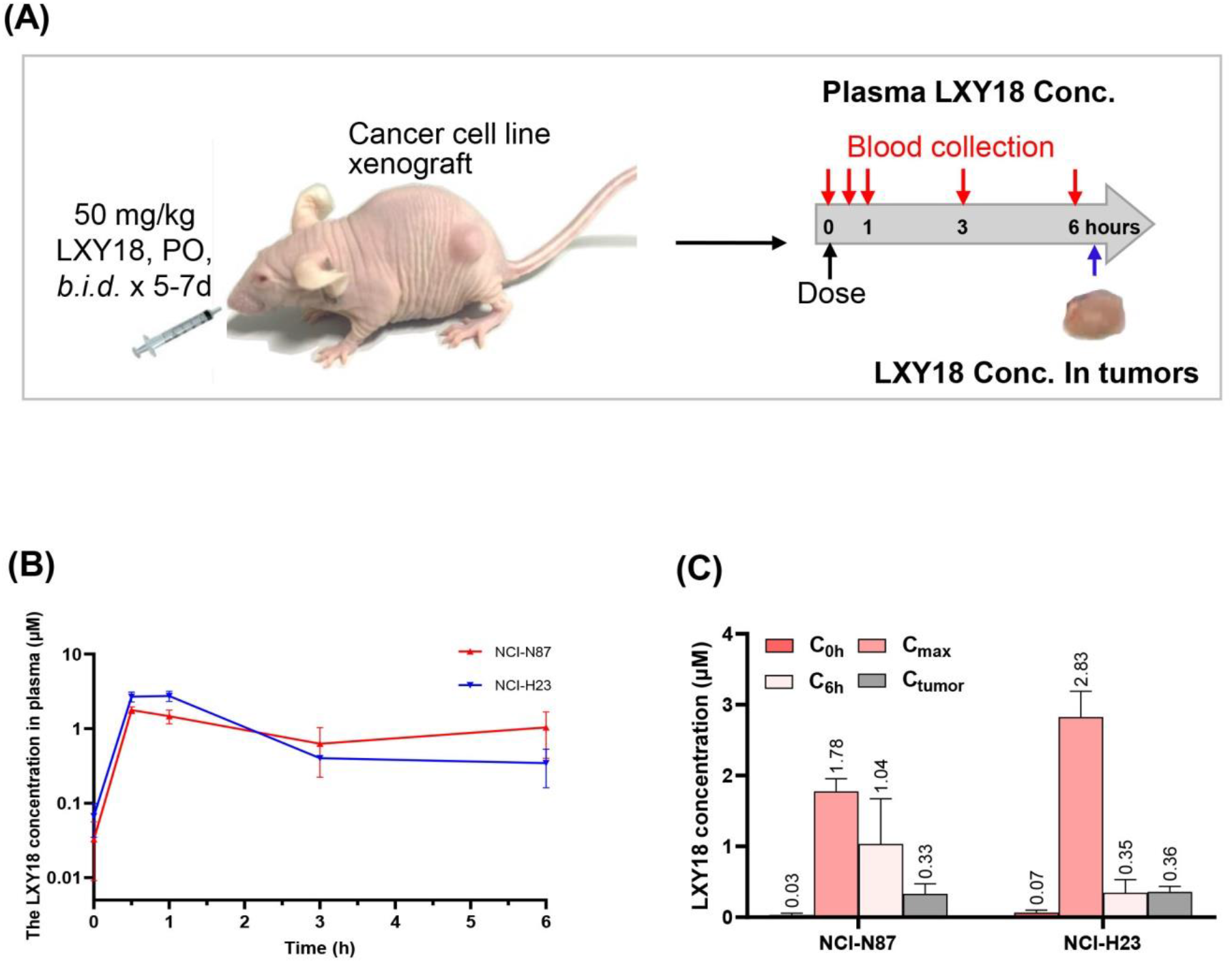
PK properties and drug concentrations in tumor tissues. Female, 12-14 weeks old BALB/c nude mice, bearing NCI-H23 and NCI-N87 xenografts were treated by oral gavage with 50 mg/kg of **LXY18** dissolved in corn oil, twice a day at 9:00 a.m. and 5:00 p.m. for five consecutive days. On the sixth day, 100 μL of blood was collected at 0, 0.5, 1, 3, and 6 h after oral administration of **LXY18** at 9:00 am. Tumors were also harvested at the last blood collection time to quantify **LXY18** concentration. Three mice were used in each group. (A) Experiment design in tumor-bearing mice to examine plasma **LXY18** exposure levels and accumulation in tumor tissues. (B) Plasma **LXY18** exposure after repeated administration. Each point, here and in C, represents the mean ± SD. (C) The **LXY18** concentration in blood and tumor tissues at 6 hours after drug administration.

Both tumor models yielded similar plasma drug concentration vs time curves (**Figure 5B**). Multiple doses to the xenografted mice decreased C_max_ 2.4-3.8 fold relative to mice that received a single dose (**Figure 5C vs Figure 3E**). Likewise, the normalized AUC_0-6h_ declined 1.6-1.7 fold (**Figure 5B vs Figure 3B**). The plasma concentrations of **LXY18** at 6 h were comparable among two mouse models with different xenografts (0.35 ± 0.18 μM for NCI-H23 versus 1.04 ± 0.64 μM for NCI-N87) (**Figure 5B**). **LXY18** concentrations in the corresponding tumor tissues approximated those found in the plasma (**Figure 5C**).

The unbound fraction, not the total amount of a drug is responsible for its pharmacological and/or toxic effects because the free form of a drug is the principal determinant of tissue distribution. **LXY18** displayed a moderate protein binding rate of 61.1% in human serum and a higher protein binding rate of 90.1% in mouse plasma (**Table S5**). After correcting for the fraction bound to plasma protein, free **LXY18** concentrations were still much higher than their corresponding MEC values, indicating that persistent disablement of AURKB could be achieved in repeated treatment.

Given the favorable oral bioavailability and accumulation in tumor cells observed *in vivo*, **LXY18** was deemed suitable to test for efficacy in murine tumor models. We chose the aggressive gastric carcinoma NCI-N87 and lung cancer cells NCI-H23, both of which responded well to **LXY18** *in vitro*. Mice bearing NCI-N87 were treated with **LXY18** whereas the control group received an equal volume of vehicle (corn oil). No obvious adverse effects were observed in the treatment group and body weight remained consistent (**Figure 6A**). **LXY18** significantly reduced tumor volume, with a TGI of 55.13% ± 29.05% (*p < 0*.*001*) (**Figure 6B**), and tumor weight at the endpoint (**Figure 6C, D**). Consistent with the observed reduction in xenograft volume, the percentage of cells positive for Ki-67 in the treatment group (33.94% ± 10.06%) was significantly reduced relative to the control group (59.4% ± 6.34%, *p=0*.*0004* by Student’s t-test) (**Figure 6E, F**). The blood vessel density in **LXY18**-treated tumors was reduced relative to the size-matched tumors in the control group (**Figure S5**). **LXY18** showed no sign of suppressing xenografts of NCI-H23 in NCG mice (**Figure S6**), indicating the importance of identifying predictive biomarkers for LXY18.

**Figure 6.**
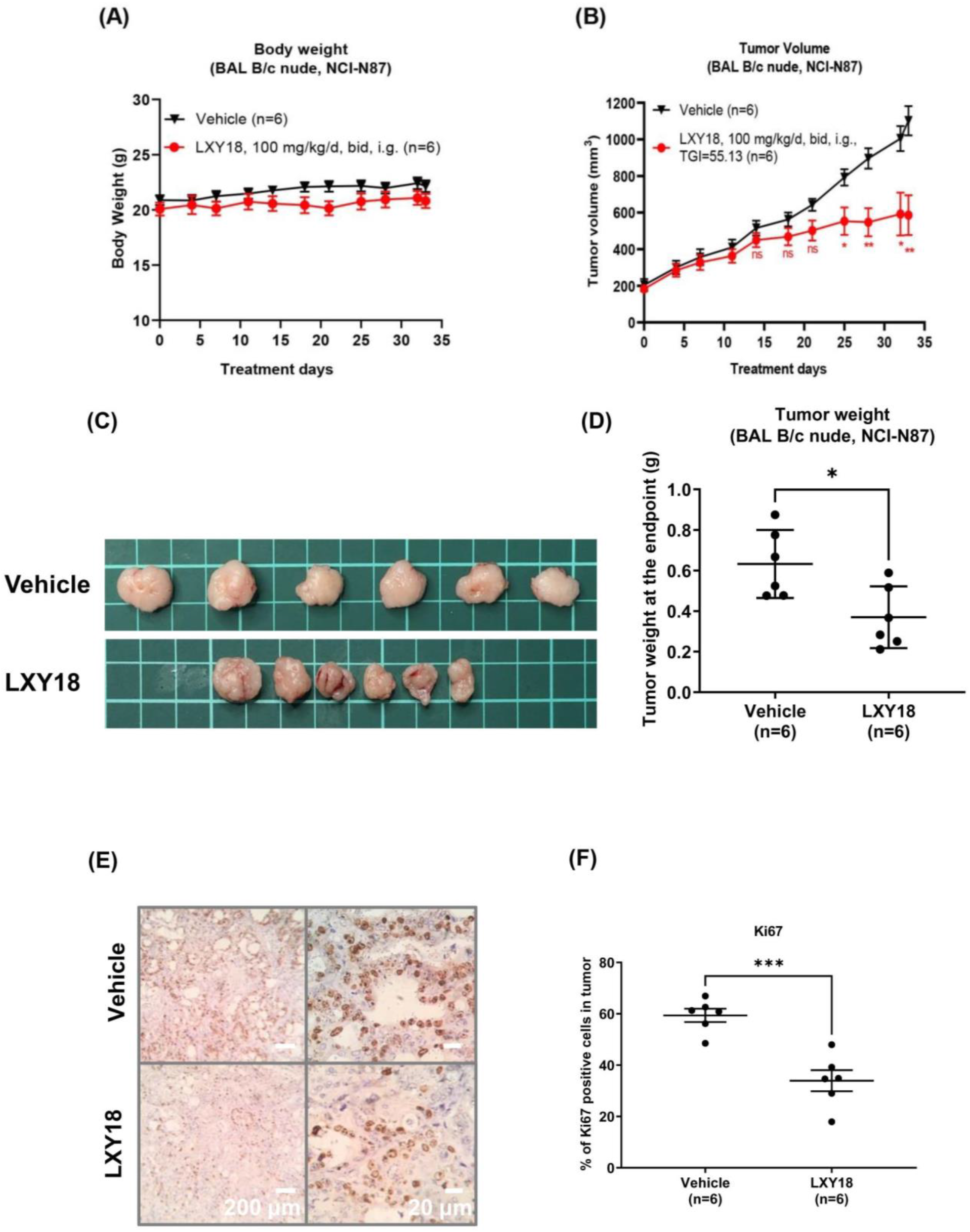
**LXY18** inhibits the growth of NCI-N87 human gastric carcinoma xenografts. Mice-bearing xenografts were treated with **LXY18** for 33 days (100 mg/kg b.i.d.), monitored twice a week for tumor sizes, and then euthanized to harvest tumor tissues. (A) Mouse body weight. (B) Tumor growth curve. (C) Gross morphology of Tumors. (D) Tumor weight. (E-F) Tumor proliferation. Tumor tissues were immunohistochemically stained for the proliferation marker Ki67 (E). Three randomly chosen fields in each tumor section were scored for Ki67 positive cells and data from n=6 mice are presented as mean ± SEM (F). *p-values* were calculated using an unpaired Student’s t-test using GraphPad Prism 8.0.2 software. (ns=not significant: *<0.05%, **<0.01%,***p<0.001%).

## Discussion and conclusions

Quinoline-based derivatives have been developed that inhibit the catalytic activity of several mitotic kinases, including AURKs.^15^ We explored non-catalytic inhibitory compounds from this class, which work by mislocalizing AURKB and other CPP complex components.^7^ Here, we have further characterized this activity for an optimized group of halogenated compounds, of which Br-substituted **LXY18** and other heavy halogen-substituted analogs are shown to act as potent AURKB relocation blockers.

**LXY18** can be delivered as an effective oral oncology agent, with a potent and broad spectrum of antimitotic activity in cancer cell lines, widespread tissue distribution, and the ability to cross the blood-brain barrier. The first generation of quinoline-based AURKB mislocalizers had limited activity *in vivo* and fail to suppress tumorigenesis in our recently published study.^7^ In the present study, **LXY18** becomes the first reported quinoline-based compound that suppresses tumorigenesis by preventing the mitotic relocation of AURKB.

**LXY18** might prevent the AURKB repositioning by binding and interfering with any component of the CPP complex. Furthermore, a variety of mitotic regulators are essential for the timely positioning of AURKB at the spindle midzone.^16^ Disabling any of these proteins or blocking their interaction with the CPP complex might hinder the relocation of AURKB at anaphase onset. Testing the physical interaction between **LXY18** and these regulators individually in a label-free assay might provide insight into the mode of action of **LXY18**. Alternatively, pulldown assays followed by LC-MS/MS analysis of bound protein might reveal a mitotic target for **LXY18**.

In summary, we have found that **LXY18** possesses potent *in vitro* and *in vivo* anticancer activity that can be delivered effectively using an oral formulation. Further characterization of metabolism, toxicity profile, and relevant target of **LXY18** and the identification of biomarkers that predict treatment response will support continued advancement toward clinical applications.

## Safety Statement

No unexpected or unusually high safety hazards were encountered. **Associated content**

## Supporting information

The supporting PDF file includes additional figures and tables of bioassays, analytic data(^1^H NMR), general experimental procedure, starting material preparation, and biology experimental methods.

## Author contributions

G.L. synthesized all compounds. Q.S., C.L.Y., J.K., Y.L., and H.M.L. performed all biology experiments. T.Z. and X.H.Z. performed in vivo PD/PK experiments. J.H.L. performed HPLC and LC-MS/MS analyses. Q.S., T.A., N.C, J.Z., S.Q.Z., and D.Y. participated in the biological data analysis. G.L., J.H.L., and N.N. analyzed NMR and LC-MS/MS data. All participated in writing the manuscript. D.Y. finalized the manuscript. All authors are employed by the J. Michael Bishop Institute of Cancer Research and/or Anticancer Bioscience, Chengdu, China

## Acknowledgment

We thank all members of the DMPK team, in particular. Drs. Manikandan and Kishore for their critical review of this manuscript and constructive suggestions.

## Abbreviations used

AURKA: Aurora Kinase A
AURKB: Aurora Kinase B
AURKB-SM: AURKB-spindle midzone
AURKB-CA: Catalytic activity of AURKB
BAR: Blockers of AURKB relocation
cLogP: Calculated octanol/water partition coefficient
CPP: Chromosomal passenger protein
DAPI: 4′,6-diamidino-2-phenylindole
DMF: N,N dimethylformamide
DMEM: Dulbecco’s Modified Eagle Medium
DCM: Dichloromethane
DIPEA: N,N-diisopropylethylamine
DMF: N,N-dimethylformamide
DMAP: 4-dimethylamino pyridine
MEC: Minimum effective concentration
MIPS: Mechanism-informed phenotypic screening
NaH: Sodium hydride
OD: Optical density
PE: Petroleum Ether
RPE: Retinal pigment epithelium
rt: Room temperature
SARs: Structure-activity relationships
THF: Tetrahydrofuran
TLC: Thin Layer Chromatography

## Availability of data and materials statement

All data analyzed during this study are included within the manuscript (and its Supplementary Information files). Data deposition does not apply to the current study.

