## Supplementary information file for "An orally bioavailable 4-phenoxy-quinoline compound as a potent AURKB relocation blocker for cancer treatment"

### Table of contents


**1.Plasma stability and physicochemical properties**

**2.Bioassays**

2.1. Cell culture

2.2. Mechanism-informed phenotypic screening assay

2.3. Immunofluorescent Staining

***3. In vitro* studies**

3.1. Determination of *in vitro* DMPK Properties

3.2. *In vitro* Stability in Plasma and Serum

3.3. pH Stability and Solubility

**4. Calculation of Lipinski and pharmacokinetic parameters**

**5. Octanol-PBS Partition Coefficient (Log D)**

**6. Plasma and Serum Protein Binding**

6.1. Determination of the Drug Concentration in Plasma and Tissues

6.2. Sample Preparation and Processing

**7. Preclinical formulation of LXY18**

**8. Animal experiments**

8.1. Rapid PK in mice and full PK in rats

8.2. Drug tissue distribution in rats

8.3. PK profiles after repeat dosing in mouse tumor models

8.4. Therapeutic efficacy in mouse tumor mice

8.5. Immunohistochemistry of tumor tissues

**9. Supporting Information Tables**

**10. Supporting Information Figures**

**11. CHEMISTRY EXPERIMENTAL SECTION**

11.1. General Methods

11.2. Chemical synthesis and Analytical Data of Compounds

11.3. Spectral Data of Compounds (figures)

**12. References**

#### 1. Plasma stability and physicochemical properties

The presence of an acetamide group in 4-phenoxyquinoline derivatives raises the issue of metabolic instability in mammalian plasma rich in amidase and an acidic environment.[1] Stability tests were performed in human serum, mouse plasma, or a buffer with a pH value of 2.2 to mimic the stomach acidity or 7.4, the physiological pH of plasma. There were no detectable levels of any degradation product, and the amount of these analogs was constant during 2-h incubation (**Tables S2** and **S3**).

They were partially soluble at $pH_{7.4}$, with the unsubstituted compound LG178 displaying the highest solubility of 61.0 ± 7.9 μM (**Table S3**). R1-substitutions reduced solubility to a range from 8.3 ± 2.9 μM for SY142 to 16.0 ± 5.7 μM for LG182. Attributable to the presence of a shared ionizable quinoline group, their solubility was enhanced by more than 10-fold at $pH_{2.2}$ relative to $pH_{7.4}$, reaching a range from 169.2 ± 6.8 μM (LXY18) to 191.0 ± 4.9 μM (LG182) compared with 200.6 ± 5.9 μM for the unsubstituted compound LG178. The experimentally determined Log D values of all six compounds were also reduced under the acidic pH relative to a neutral pH (**Table S3**). The much greater reduction of Log D values was seen with LG178 and its OMe-substituted derivative relative to their halogen-contained analogs. Nevertheless, log D values for all compounds at either of the two pH values satisfied the rule of five for oral drugs.[2]

#### 2. Bioassays

#### 2.1. Cell culture

The human retinal pigment epithelial cell lines RPE-MYC$^{H2B\text{-}GFP}$ and RPE-MYC$^{Bcl2}$ were described.[3] Lung cancer cell lines (NCI-H460, NCI-H23, NCI-H596, NCI-H841, A549, and NCI-H2170), colon cancer cell lines (SW480, HCT-116, and HCT-116 p53-/-), breast cancer cell lines (MCF-7, MDA-MB-175IV, MDA-MB-435), gastric cancer cell lines (NCI-N87 and HTB135), skin cancer cell lines (UACC-62 and C32), a prostate cancer cell line DU145, an ovary cancer cell line Caov-3 and a cervical cancer cell line Hela were obtained from American Type Culture Collection (ATCC). All lines were passaged in DMEM (Gibco, Cat. No. 12100061) or RPMI-1640 (Gibco, Cat. No. C11875500BT) supplemented with 10% fetal bovine serum (Excell, Cat. No. FSP500 10099141), penicillin (100 U/mL)-streptomycin (100 μg/mL) (Gibco, Cat. No.15140-122), 2 mM L-glutamine (Gibco, Cat. No. 25030081), and 1 mM sodium pyruvate (Gibco, Cat. No. 11360070). Cells were maintained at 37 °C with 5% $CO_2$ in a humidified incubator. Mycoplasma contamination of cell culture was evaluated by using a Myco-LumiTM Luminescent Mycoplasma Detection Kit (Beyotime, Cat. No. C0298M) before experiments were initiated and after the experiments were finished.

#### 2.2. Mechanism-informed phenotypic screening assay

An image-based screening assay of live cells to identify compounds that phenocopy the genetic loss of the chromosomal passenger protein complex was described.[4] Briefly, a screening cell line was passaged into batches of 96-well plates 18-24 h before exposure for 72 h to a test compound at concentrations between 0.5 nM and 1 μM with a 2-fold serial dilution. The DNA was visualized with H2B-GFP in live cells or stained with DAPI in fixed samples to determine the minimally effective concentrations that elicit multinucleation, mitotic arrest, and cell death. Cells were analyzed by a fluorescent cell imager (GE IN-Cell Analyzer 2000) or an EVOS FL Auto microscope. Cells treated with 0.1% of DMSO were used as the control for the assay.

##### 2.3. Immunofluorescent Staining

Immunofluorescent analysis was performed as described[3] without modification after cells were cultured overnight on glass coverslips (Φ14mm) pre-coated with 0.1% gelatin in a six-well plate. Cells were counterstained with DAPI (Yeasen, Cat. No. 36308ES20) in the mounting media. Images were acquired under an EVOS auto FL Imaging System (ThermoFisher). The immunofluorescent signal intensity was quantified with the ImageJ software.

**Antibody information:** The primary antibodies used here were a rabbit anti-AURKA Thr288P antibody (Cat. No. 3079) and a rabbit anti-H3Ser10P (Cat. No. 53348) from Cell Signaling Technology. To examine the mitotic localization of AURKB, a rabbit anti-AURKB antibody (Cat. No. abs131460) from Absin (Shanghai, China) was used. Human autoimmune serum CREST which recognizes a variety of kinetochore proteins was a gift from B. R. Brinkley (Baylor College of Medicine, Houston). Secondary antibodies including Rhodamine (TRITC) AffiniPure Goat Anti-Rabbit IgG (H+L) (Cat. #111-025-003), Rhodamine RedTM-X-conjugated AffiniPure Donkey Anti-Human IgG (H+L) (Cat. No. 709-295-149) and Fluorescein (FITC)-conjugated Affinipure Goat Anti-Rabbit IgG(H+L) (Cat. No. 111-095-003) were from Jackson ImmunoResearch.

#### 3. *In vitro* studies

##### 3.1. Determination of *in vitro* DMPK Properties

LC-MS Conditions. *In vitro* DMPK properties were determined with an LC-MS system (Agilent InfinityLab LC/MSD iQ) with the same instrument conditions as described.[5]

##### 3.2. *In vitro* Stability in Plasma and Serum

Human serum AB (Cat#100512) and mouse plasma (Cat#SND-X0102) were from Geminibio (California, USA) and Shinuoda (Chuzhou, China), respectively. The stability of LXY18 and its analogs was studied at a concentration of 1 or 10 μM. The samples were prepared by spiking mouse plasma or human serum with a test compound diluted from a stock solution with acetonitrile in 1.5 mL Eppendorf tubes and then incubated for 2 h at 37.5 °C in a water bath. 40 μL of the incubated samples were then mixed with 460 μL acetonitrile by vortexing (3000 rpm, 2 min) and the mixture was then centrifuged at 14000 rpm for 10 min. 150 μL of the supernatant in each Eppendorf tube was transferred into a liner pipeline of 1.5 mL vials for analysis by LC-MS.

##### 3.3. pH Stability and Solubility

An aliquot of 10 μL of a test compound was diluted with 990 μL of an HCl buffer ($pH_{2.2}$) or a 10 mM PBS buffer ($pH_{7.4}$) in 1.5 mL Eppendorf tubes to obtain 200, 100, and 50 μM samples. The tubes were placed at room temperature for 0, 2, 24, and 72 h and then aliquots of solution were taken to quantify the compound and detect its degradation products by LC-MS.

**4. Calculation of Lipinski and pharmacokinetic parameters**

Lipinski Parameters were performed by using the online open-source cheminformatics.[6] The pharmacokinetic parameters of the compounds were accomplished by the SwissADME tool.[7]

**5. Octanol-PBS Partition Coefficient (Log D)**

Log D was determined at pH 2.2 and pH 7.4 as we described before without modification.[5]

**6. Plasma and Serum Protein Binding**

Plasma protein binding was performed by the equilibrium dialysis method with slight modification from published literature.[8] The samples (10 μM of LXY18) were prepared by spiking blank mouse plasma and human serum with LXY18 diluted from a stock solution with acetonitrile and placed in a water bath (2 h, 37.5 °C) for thorough binding interaction. The mixture (200 μL) in a snakeskin dialysis tube (Thermo Scientific, 10 K MWCO) was then dialyzed for 18 h at 37°C against 40 mL of dialysis buffer (0.01M PBS, pH 7.4) Finally, the dialysis buffer was taken to quantify LXY18 by LC-MS. The plasma/serum protein binding rate was calculated by employing the following equation:

Binding rate (%) = ($Amount_{200\ \mu L\ mixed\ samples}$ – $Amount_{dialysis\ buffer}$) / $Amount_{200\ \mu L\ mixed\ samples}$ × 100%

##### 6.1. Determination of the Drug Concentration in Plasma and Tissues

LC-MS/MS Conditions. The LXY18 concentrations in plasma and tumor tissues were determined using a Thermo Scientific Vanquish UHPLC system coupled with a TSQ Quantis™ Triple Quadrupole Mass Spectrometer. Except for MS/MS scan mode setting, LC-MS/MS conditions are the same as in the previous report.[9] Selective reaction monitoring (SRM) is performed on a scanning mass spectrometer. SMR parameters are set according to the precursor(m/z), product (m/z), collision energy (V), and RF LENS (V) of each compound and were set as follows:

| Compound | Q1 (m/z) | Q3 (m/z) | Collision energy (V) | RF lens (V) |
|---|---|---|---|---|
| LG178 | 309.17 | 267.1 | 23 | 151.2 |
| LXY19 | 327.2 | 285.0 | 24 | 164.2 |
| LG182 | 339.2 | 297.2 | 26 | 126.0 |
| SY142 | 343.1 | 301.0 | 24 | 168.2 |
| LXY18 | 389.1 | 347.1 | 25 | 153.5 |
| LC08 | 435.1 | 393.0 | 25 | 190.4 |

##### 6.2. Sample Preparation and Processing

Plasma samples were prepared and processed as previously described.[9] The extracted solutions were analyzed by the LC-MS/MS system mentioned above.

**7. Preclinical formulation of LXY18**

1. **HPMC formulation**: Solid powder of LXY18 was added to a solution of 0.5% HPMC (pH2.2), and homogenized for 30 min by using an ultrasonic cell disruptor (Ning Bo Xin Zhi, JY92-IIN), resulting in a milky white uniform suspension of LXY18 at a concentration of 10 mg/mL. The formulation was stored at 4℃.

2. ***PEG 300 formulation:*** Solid powder of LXY18 was added in DMSO and fully dissolved by vortexing followed by sonication in a water bath ultrasonic (BRANSON, 1510) for 10 min. The solution was diluted at 1:9 (v/v) with a PEG 300 solution that contains 50% v/v PEG 300 and 1% v/v Tween 80 in 1x PBS buffer at pH 8.0. The mixture was then blended by vortexing and then sonicated in a water bath ultrasonic for 10 min to yield a clear solution of LXY18 at a concentration of 10 mg/mL. The formulation was stored at 4℃.

3. ***Corn oil formulation:*** Solid powder of LXY18 was first added to corn oil and then homogenized for 30 min by using an ultrasonic cell disruptor to produce a yellow suspension with LXY18 at a concentration of 10 mg/mL. The formulation was stored at 4℃.

**8. Animal experiment:**

*Ethics statement*

The study was conducted according to the animal study protocols with reference number (**BUA#: BU000021-010**) and was approved by the institutional animal care and use committee (IACUC guideline 20.20) of the Institutional Biosafety Committee (IBC) (or ethical committee) at MBICR**.** Directions for $CO_2$ euthanasia of Rodents (mice) were done under (IACUC guideline 20.20). This study was carried out in compliance with the ARRIVE guidelines.

*Mouse information*

Six weeks old female BAL B/c nude mice were sourced from GemPharmatech experimental animals Co., LTD to conduct animal experiments.

##### 8.1. Rapid PK in mice and full PK in rats

PK studies followed the protocols describe before. [5,10] Briefly, Two 8-12 week-old female FVB/N mice, 20-30 g each, were used for each formulation. Three 8-week-old female Wistar rats, 180-220 g each, were used for full PK studies and dosed at 2 mg/kg of LXY18 dissolved into 50% PEG 300 at pH 5.63 by tail vein injection. Alternatively, Wistar rats were orally dosed with 2, 10, or 50 mg/kg of LXY18 suspended in corn oil, and three rats were used for each dose. Blood was collected into heparinized tubes and analyzed for the plasma concentration of LXY18 with LC-MS/MS. The area under the plasma drug concentration-time curve was calculated by the non-compartmental analysis method in Phoenix™ WinNonlin® software.

**8.2. Drug tissue distribution in rats**

Nine 9-week-old female Wistar rats, 200-240 g each, were orally dosed with 50 mg/kg of LXY18 suspended in corn oil. Animals were starved for 16 h before treatment, and food was allowed 2 h after drug administration. Free water intake was permitted throughout the study. At 1, 4, and 8 h after drug administration, body fluids and various organs were collected, including blood, heart, lung, liver, pancreas, spleen, ovary, kidney, muscle of the leg, Lymph nodes, thighbone, adipose tissue, and brain. The plasma concentration of LXY18 was analyzed with LC-MS/MS. Tissues and organs were removed from residual blood by washing with 1 x PBS. About 200 mg of tissues were then mixed with 800 μL of grinding fluid (methanol: PBS = 1:1), homogenized by using a high throughput tissue grinding machine, and finally centrifuged at 12000 rpm for 5 min to obtain the supernatant for quantification of LXY18 with LC-MS/MS.

**8.3. PK profiles after repeat dosing in mouse tumor models**

Female BAL B/c nude mice at an age of 12-14 weeks old, 19-24 g each, were used to generate xenografts of human cancer cell lines (NCI-H23 and NCI-N87). Three mice were utilized for each tumor model. Tumor-bearing mice were dosed with 50 mg/kg of LXY18 in corn oil by oral gavage twice a day for five consecutive days. A PK test was performed at 0, 0.5, 1, 3, and 6 h after the last dose. Mice were treated and blood was collected as described before.[10] Six hours after the last drug administration, mice were sacrificed, and tumors were harvested. About 200 mg of tumor tissues were mixed with 800 μL of grinding fluid (methanol: PBS = 1:1), homogenized with a high throughput tissue grinding machine, and then centrifuged at 12000 rpm for 5 min to obtain the supernatant for quantification of LXY18 with LC-MS/MS.

##### 8.4. Therapeutic efficacy in mouse tumor models

Xenografts were created by subcutaneously transplanting 1.68 million NCI-N87 cells or 2.4 million Calu-6 cells in 100 uL of PBS supplemented with 50% Matrigel into each dorsal flank of a 6-10 weeks old female immunocompromised BAL B/c nude mouse with a body weight of 19-24 g each. Tumors were allowed to grow to 6-7 mm in diameter before tumor-bearing mice were randomized into the control group (n=6) or experimental group (n=6) by using a computer program in R (version 2.0.0). The mice bearing NCI-N87 xenografts in the experimental group received a daily dose of 50 mg/kg of LXY18 by oral gavage twice a day for thirty-three consecutive days. The cohort bearing Calu-6 cancer cells received an oral dose of 80 mg/kg/day of LXY18 once a day for twenty consecutive days. For the control group, the same amount of vehicle (corn oil) was administered. Mouse body weight and tumor burden were measured as indicated in figure legends.

All data were plotted by using Graphpad Prism 8.0.2, and tumor growth inhibition (TGI) was calculated by the formula: TGI =(1-△V treatment / △V Control) x100.

##### 8.5. Immunohistochemistry of tumor tissues

At the endpoint, the mice were euthanized to harvest tumor tissues, which were fixed in 4% formaldehyde in a 0.1 M phosphate buffer (pH 7.4), embedded in paraffin, and cut into sections of 2 μm in thickness using a Leica-2016 Rotary Slicer. The sections were analyzed by immunohistochemical staining for Ki67 or CD31. the sections were treated in a microwave for 20 min in a Tris-EDTA buffer (10 mM Tris-base, 1 mM EDTA solution, 0.05% Tween 20, pH 9.0) for antigen retrieval and were then allowed to cool to room temperature over a period of 30 min before the endogenous peroxidase was inactivated with 3% $H_2O_2$ for 10 min. Incubation with the primary antibody was performed at 4℃ overnight. After washed in TBST three times, the slides were incubated with an HRP-conjugated goat anti-rabbit secondary antibody. Finally, staining was developed using an SP Kit before slides were counterstained in hematoxylin, dehydrated, and mounted. Images of slides were acquired by using a Panoramic 250, 3DHISTECH（Hungary) digital slide scanner. The presence of the proteins was scored by two blinded observers.

#### 9. Supporting Information Tables

**Table S1.** Calculated Lipinski and drug-likeness parameters of **LXY18** and its analogs

| S.No | Comps | R group | MW | HBA | HBD | cLogP | TPSA | Nrotb |
|---|---|---|---|---|---|---|---|---|
| 1 | LC08 | I | 376.33 | 4 | 1 | 5.01 | 60.45 | 4 |
| 2 | SY142 | Cl | 342.78 | 4 | 1 | 4.65 | 60.45 | 4 |
| 3 | LG178 | H | 308.34 | 4 | 1 | 3.99 | 60.45 | 4 |
| 4 | LG182 | OMe | 338.36 | 5 | 1 | 4.00 | 69.68 | 5 |
| 5 | LXY19 | F | 326.33 | 4 | 1 | 4.13 | 60.45 | 4 |
| 6 | LXY18 | Br | 387.23 | 4 | 1 | 4.76 | 60.45 | 4 |

SNO, sampler number; Cmps, compounds; MW, molecular weight; HBA, H-bond accepter; HBD, H-bond donors; cLogP, Calculated octanol/water partition coefficient; TPSA, total polar surface area; Nrotb, the number of rotatable bonds

**Table S2.** Stability of **LXY18** and its analogs in mouse plasma and human serum

| | Test Conc. | Recovery rate | | | | | |
|---|---|---|---|---|---|---|---|
| | | LG178 | LXY19 | LG182 | SY142 | LXY18 | LC08 |
| Mouse plasma | 1 μM | 84.2% | 103.1% | 94.8% | 94.3% | 98.6% | 94.8% |
| | 10 μM | 102.5% | 96.0% | 109.0% | 104.7% | 102.2% | 101.0% |
| Human serum | 1 μM | 89.6% | 102.2% | 95.0% | 96.3% | 106.8% | 99.3% |
| | 10 μM | 103.7% | 93.6% | 109.0% | 101.0% | 98.9% | 102.7% |

LXY18 and its analogs at 1 or 10 μM were incubated with mouse plasma or human serum for 2 h at 37.5 °C to calculate the recovery rate. Control incubation (recovery rate, 100%) was included for each compound tested where the compound was incubated with 0.01 M phosphate-buffered saline (PBS). Each compound was quantified by LC-MS/MS. A  representative experiment is presented.

**Table S3.** Physicochemical properties of **LXY18** and its analogs

| Properties | Condition | LG178 | LXY19 | LG182 | SY142 | LXY18 | LC08 |
|---|---|---|---|---|---|---|---|
| pH stability | $pH_{2.2}$ | Y | Y | Y | Y | Y | Y |
| | $pH_{7.4}$ | Y | Y | Y | Y | Y | Y |

| | | | | | | | |
|---|---|---|---|---|---|---|---|
| Solubility ($\mu$M) | $pH_{2.2}$ | 200.6±5.9 | 190.5±1.0 | 191.0±4.9 | 175.8±20.5 | 169.2±6.8 | 175.0±8.7 |
| | $pH_{7.4}$ | 61.0±7.9 | 13.5±2.9 | 16.0±5.7 | 8.3±2.9 | 15.6±2.1 | 12.7±5.7 |
| Log *D* | $pH_{2.2}$ | 0.27±0.03 | 1.04±0.00 | 0.31±0.03 | 1.64±0.14 | 1.95±0.02 | 1.53±0.03 |
| | $pH_{7.4}$ | 3.79±0.06 | 3.95±0.01 | 3.81±0.11 | 3.37±0.22 | 4.82±0.07 | 1.71±0.09 |

The stability and solubility were investigated after incubation for 72 h of each compound (200 $\mu$M) at the indicated pH. Three experiments with three replicates in each were tested for each of the above assays. Data are presented as mean ± SD. Y: no degradation products were detected by LC-MS.

**Table S4. The bioactivities of LXY18, SY142, LC08**

| Group | Conc. (nM) | AURKA Kinase Activity (%)[a] | AURKB Kinase Activity (%)[b] | Localized AURKB [c] | Polyploidy (%)[d] |
|---|---|---|---|---|---|
| DMSO | / | 100 ± 1 | 100 ± 3 | Yes | <1 |
| LXY18 | 10 | 100±1 | 100±2 | No | 20±5 |
| | 20 | 100±1 | 100±1 | No | 100±0 |
| | 40 | 100±1 | 100±1 | No | 100±0 |
| | 80 | 49±1 | 77±1 | NA | 100±0 |
| | 160 | 51±1 | 67±1 | NA | 100±0 |
| SY142 | 10 | 100±1 | 100±1 | No | 87±3 |
| | 20 | 100±1 | 100±1 | No | 100±0 |
| | 40 | 100±1 | 100±1 | No | 100±0 |

| | | | | | |
|---|---|---|---|---|---|
| | 80 | 59±1 | 100±1 | No | 100±0 |
| | 160 | 48±1 | 100±1 | NA | 100±0 |
| LC08 | 5 | 100±1 | 100±0 | No | 85±4 |
| | 10 | 100±1 | 100±0 | No | 100±0 |
| | 20 | 65±1 | 100±0 | No | 100±0 |
| | 40 | 52±1 | 100±0 | NA | 100±0 |
| | 160 | 39±1 | 100±0 | NA | 100±0 |

RPE-MYC$^{Bcl2}$ cells were treated with the indicated compounds for 6 or 72 h prior to analysis for the indicated phenotypes. The cells treated with vehicle (0.1% DMSO) were used as a negative control. Data are presented as mean +/- SD from three independent experiments.

[a]AURKA kinase activity was determined by quantifying the immunofluorescent staining for AURKA Thr288P 6 h after initiation of drug treatment. [b]AURKB kinase activity was determined by quantifying the immunofluorescent staining for H3 Ser10P 6 h after initiation of drug treatment. [c] Localization of AURKB at the spindle midzone in anaphase cells was determined by immunofluorescent staining for AURKB 6 h after initiation of drug treatment. Yes, positive for localized AURKB; No, negative for localized AURKB; NA, not assayed. [d]The percentage of multinucleated polyploid cells in the fully attached interphase population was quantified 72 h after drug treatment.

**Table S5.** Plasma and serum protein binding rate of **LXY18**

| | Mouse plasma binding rate (%) | | Human serum binding rate (%) | |
|---|---|---|---|---|
| | 1 | 2 | 1 | 2 |
| Value | 91.4% | 88.8% | 59.7% | 62.4% |
| Average | 90.1% | | 61.1% | |

The plasma and serum protein binding rates of LXY18 were determined by the equilibrium dialysis at a compound concentration of 10 μM in two independent experiments. LXY18 was quantified by LC-MS/MS.

#### 10. Supporting Information Figures

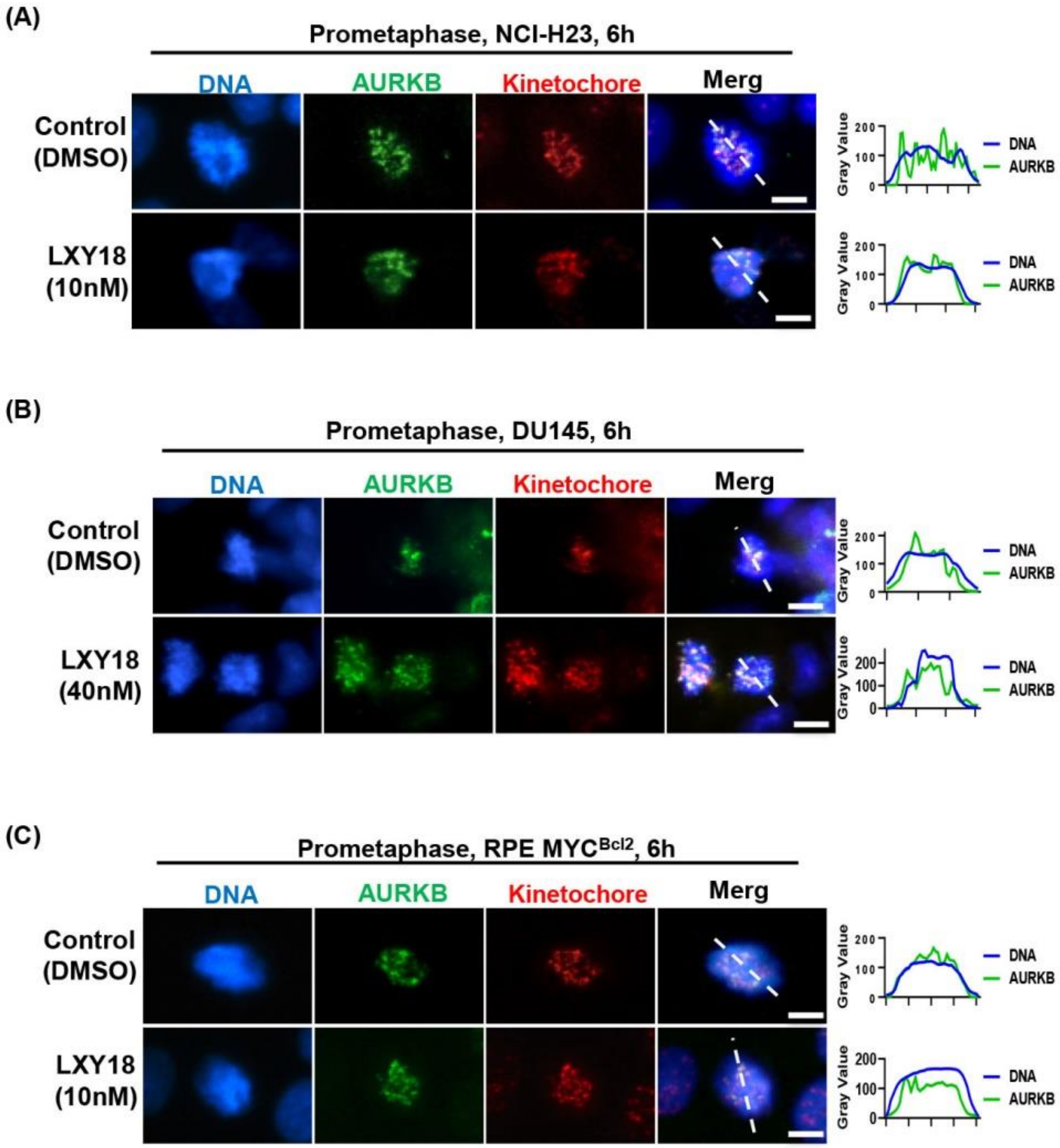

**Figure S1. LXY18** has no effect on the localization of AURKB at prometaphase kinetochores.

NCI-H23 (A), DU145 (B), and RPEMYC$^{Bcl2}$ (C) cells were treated with either vehicle control (0.1% DMSO) or LXY18 (10nM or 40 nM) for 6 h before immunofluorescent analysis of AURKB and kinetochore proteins. DNA was stained with DAPI. Kinetochore proteins were detected with a human autoimmune CREST serum. For quantitation, the fluorescence intensity along the white dashed line was determined by using ImageJ software and this was plotted as a relative gray value versus distance in pixels. Scale bars: 10 µm. For each group, more than 20 mitotic cells were scored and displayed consistent results. A representative cell image from each group is presented.

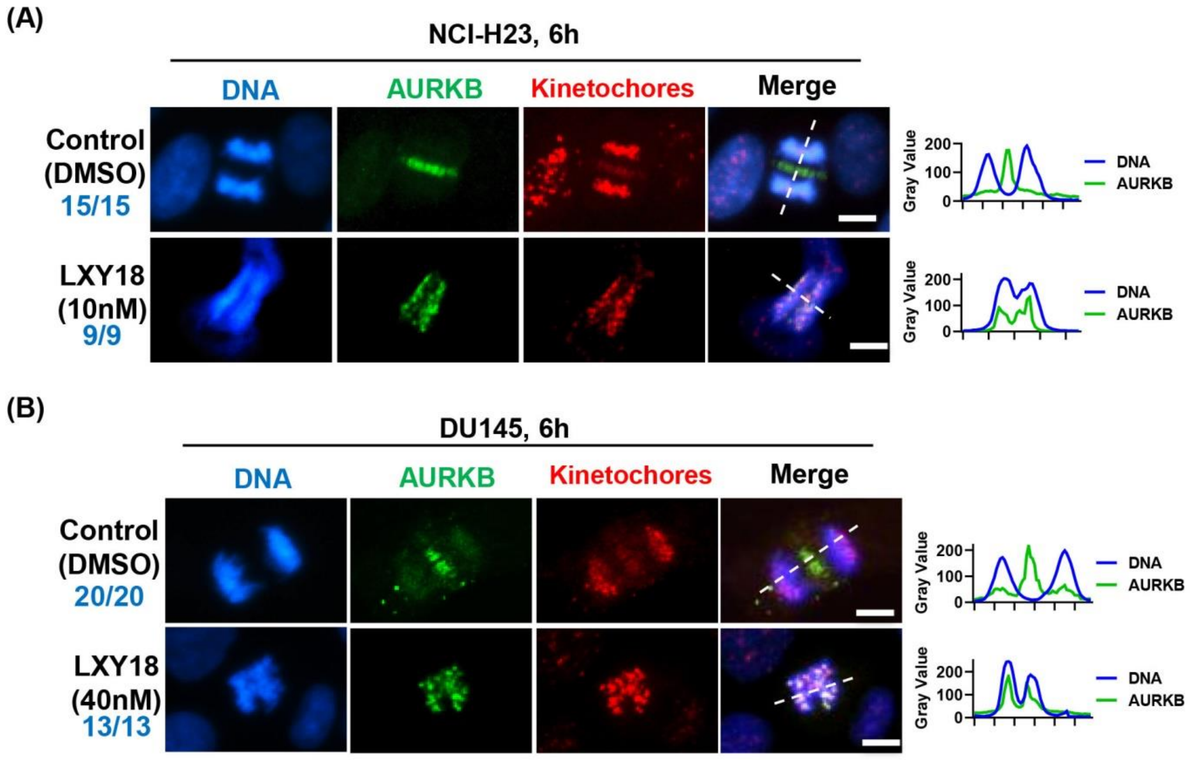

**Figure S2.** LXY18 blocks the relocation of AURKB from chromosomes to the spindle midzone on the anaphase onset.

NCI-H23 (A) and DU145 (B) cells were treated with either vehicle control (0.1% DMSO) or the indicated concentration of LXY18 for 6 h before immunofluorescent analysis of AURKB and kinetochore proteins. DNA was stained with DAPI. Kinetochore proteins were detected with a human autoimmune CREST serum. For quantitation, the fluorescence intensity along the white dashed line was determined by using ImageJ software and this was plotted as a relative gray value versus distance in pixels. Scale bars: 10 μm. The numbers in each group indicate the number of mitotic cells positive for the indicated phenotype as well as the number of mitotic cells scored.

(A) Cell line: DU145

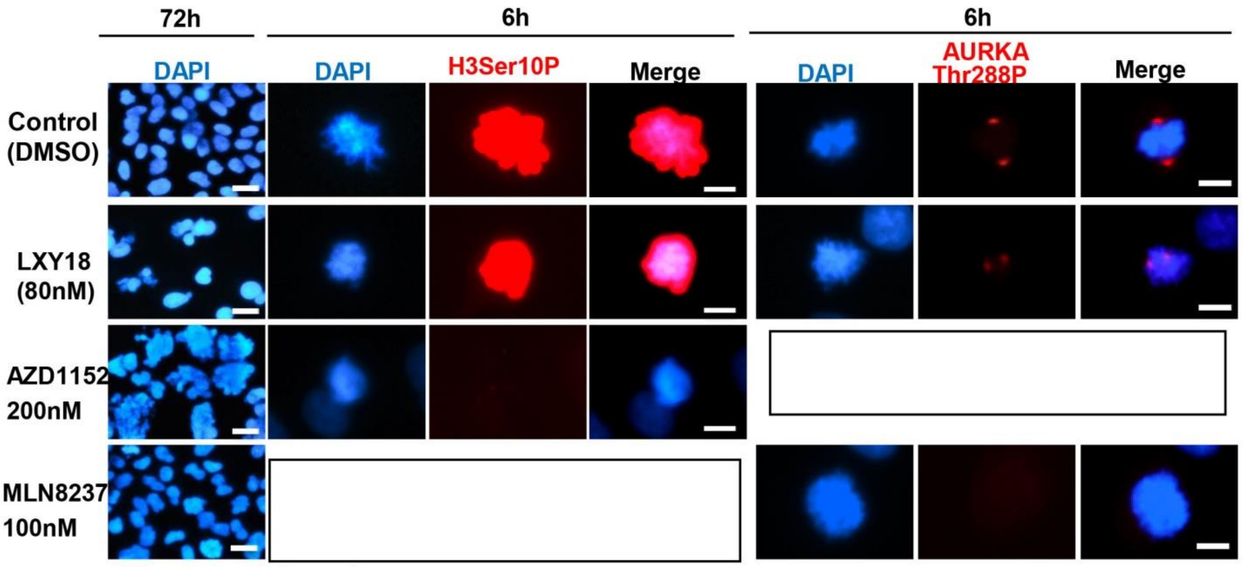


(B) Cell line: RPE MBC

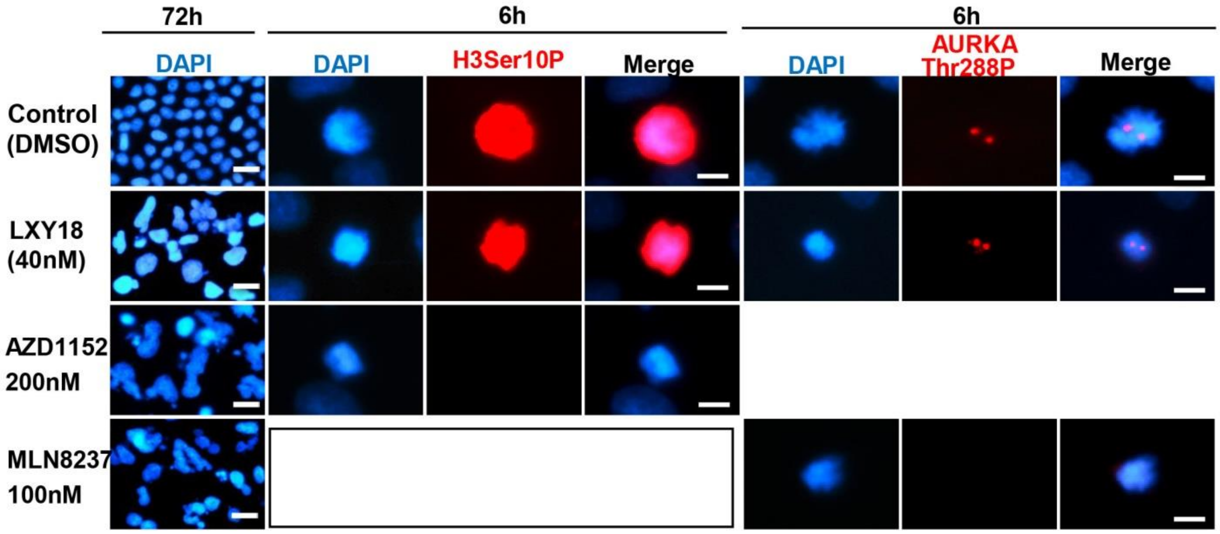


**Figure S3. LXY18 fails to inhibit the catalytic activities of AURKA and AURKB.** DU145 (A) and RPE-MYC$^{BCL2}$(B) cells were treated with indicated compounds for 6 h before immunofluorescent analysis of H3Ser10P and autophosphorylation of Threonine

288 of AURKA (AURKA-Thr288P). DNA was also stained with DAPI after 72 h of treatment to visualize polyploid nuclei. Cells treated with vehicle (0.1% of DMSO) were used as a negative control. All scale bars: 10 μm. More than 20 mitotic cells in each group were scored and all displayed similar images as the representative cell presented.

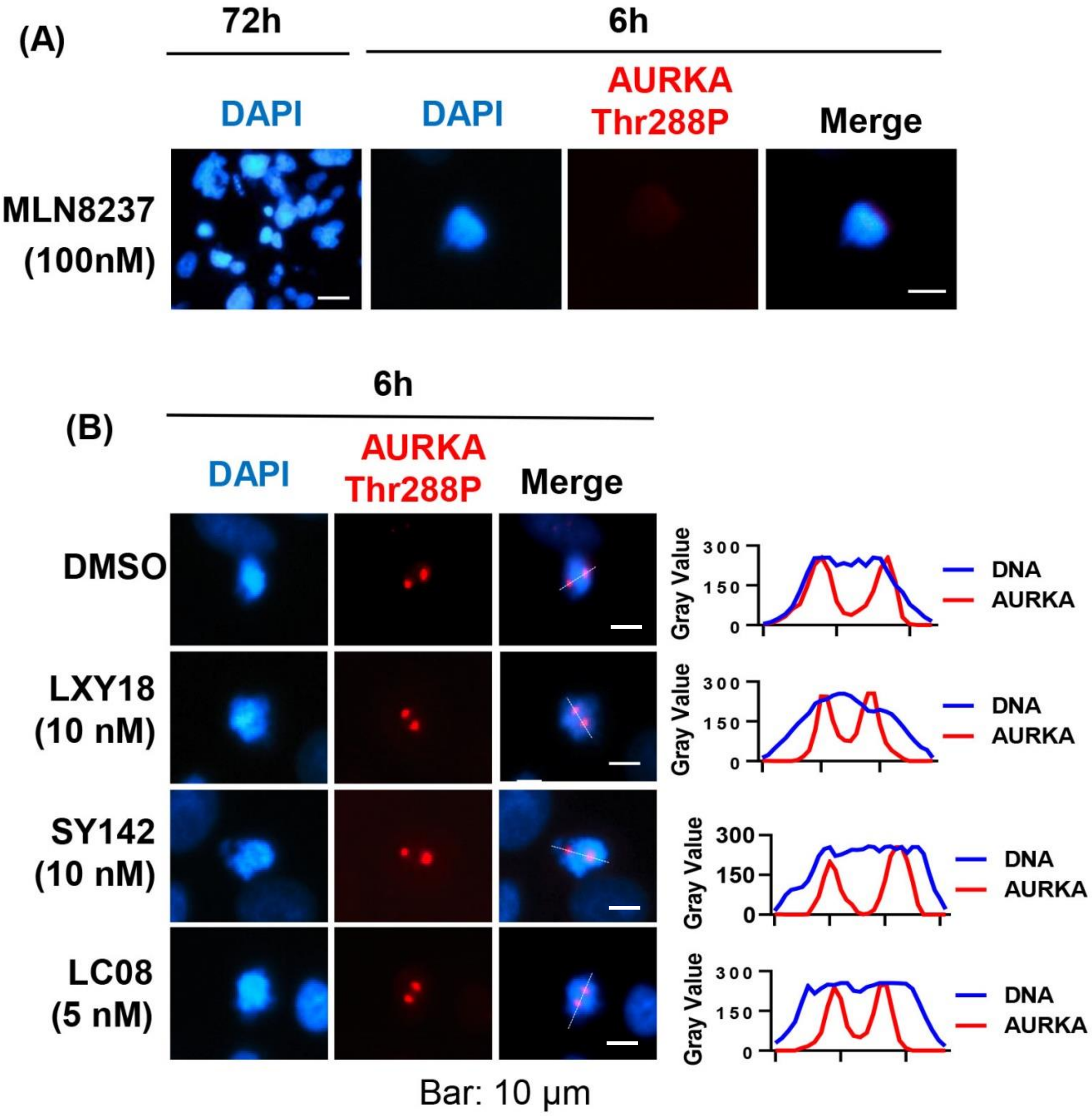


**Figure S4. LXY18 and its analogs SY142 and LC08 do not inhibit the catalytic activities of AURKA.** RPE-MYC$^{BCL2}$ cells were treated with AURKA-specific inhibitor MLN8237 (A) or LXY18 and its two analogs SY142 and LC08 (B) for 6 h before immunofluorescent analysis of autophosphorylation of Threonine 288 of AURKA

(AURKA-Thr288P). DNA was also stained with DAPI after 72 h to visualize polyploid nuclei (A, B). For quantitation in B, the fluorescence intensity along the white dashed line was determined by using ImageJ software and this was plotted as a relative gray value versus distance in pixels. Cells treated with vehicle (0.1% of DMSO) were used as a negative control. All scale bars: 10 μm. More than 20 mitotic cells in each group were scored and all displayed similar images as the representative cell presented.

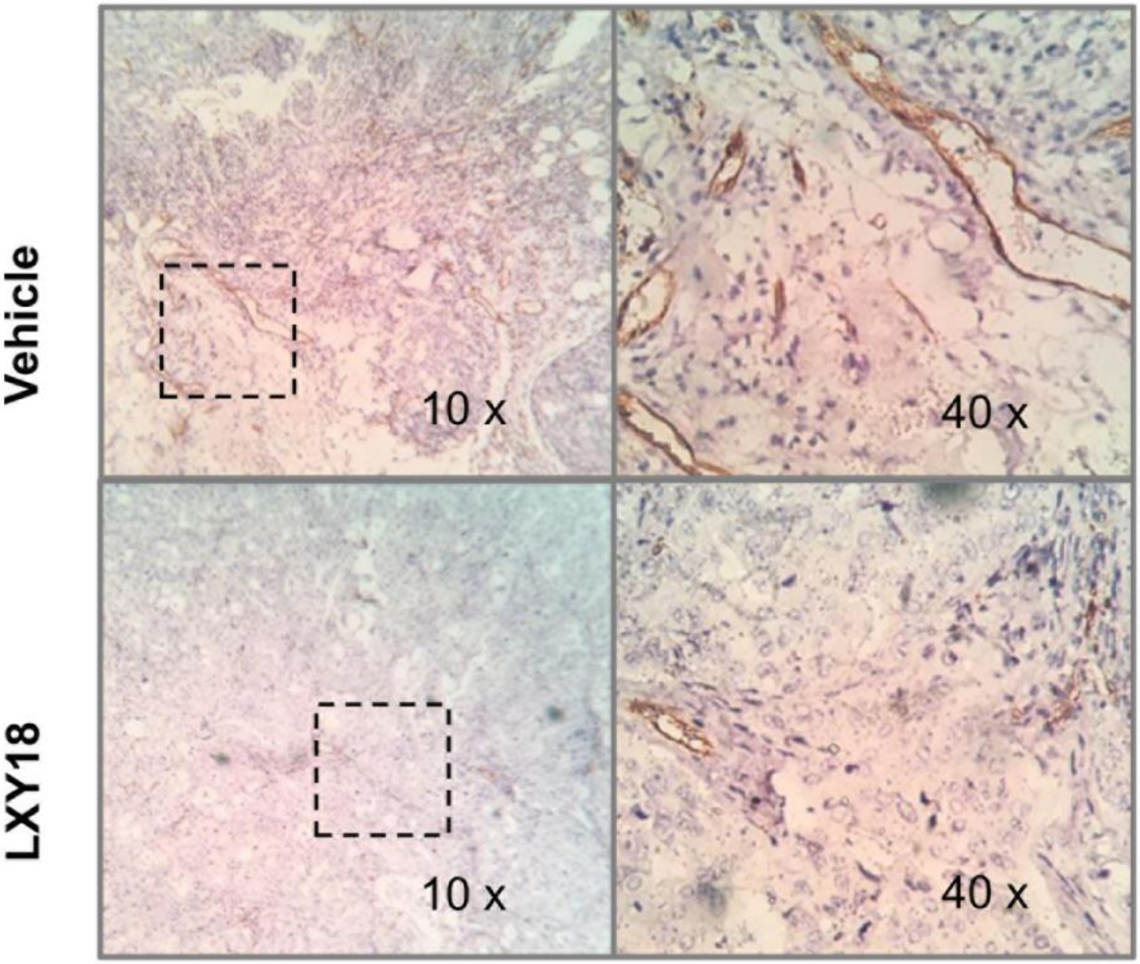


**Figure S5. LXY18 suppresses angiogenesis.** N87 xenografts described in Figure 6 were subjected to immunohistochemical staining for angiogenesis marker CD31.

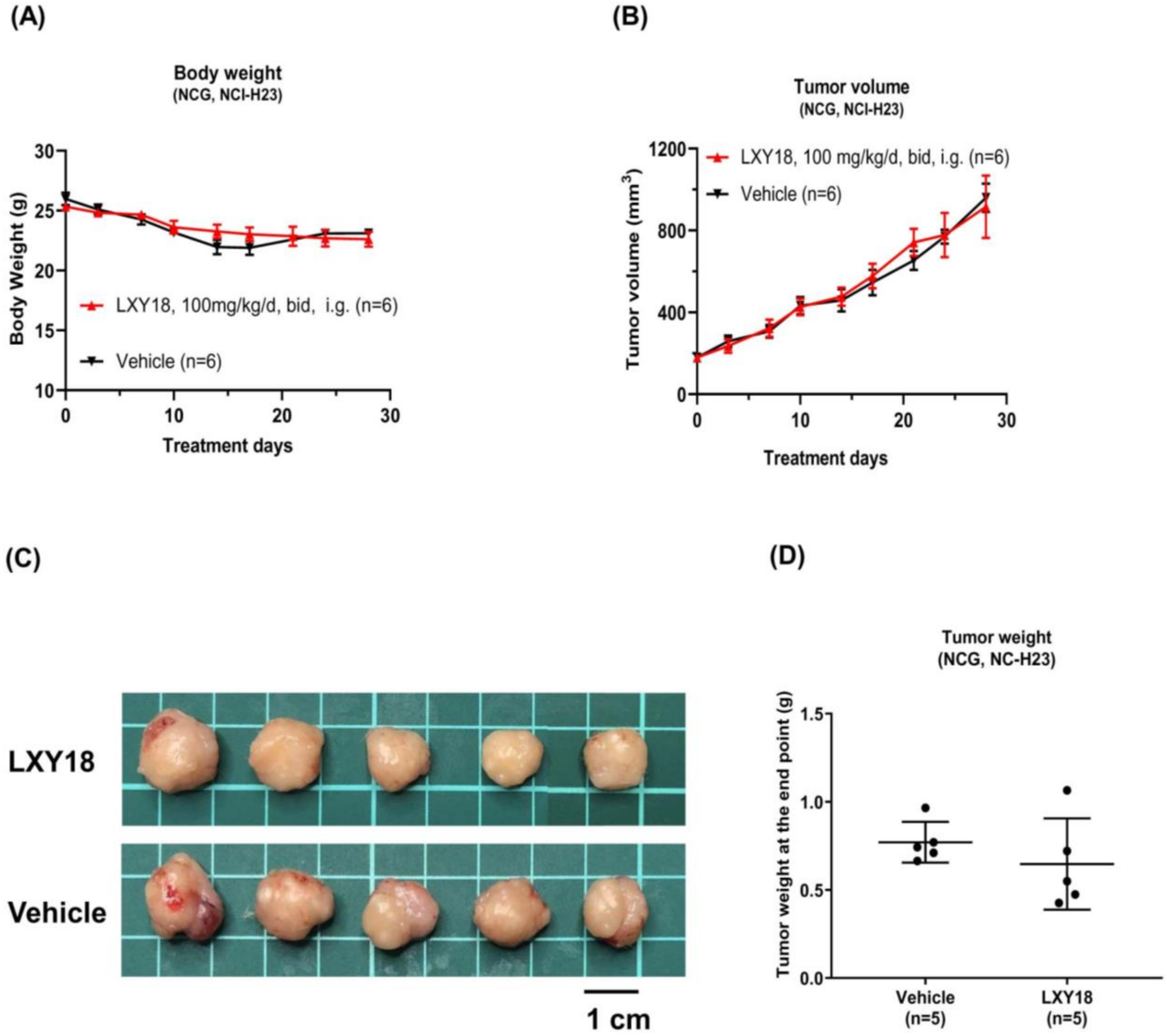


**Figure S6. LXY18 fails to suppress the growth of NCI-H23 xenograft in mice..** Six 6-week-old female NCG mice, 20-24 g each, bearing a xenograft of the NCI-H23 non-small cell lung cancer cell line. Days 0-14 were treated with 12.5 mg/kg/d of LXY18, days 15-30 were 100 mg/kg/d, twice a day, and then euthanized to harvest tumor tissues. The body weight (A) and tumor volume (B) were measured twice a week. The dissected tumors at the endpoint are shown in C and the tumor weight in D.

#### 11. CHEMISTRY EXPERIMENTAL SECTION

***11.1. General Methods***. Unless stated otherwise, all the chemicals required for synthesis were purchased from commercially available suppliers and used without further purification. $^{1}$H NMR and $^{13}$C NMR spectra of the synthesized compounds were recorded on a Bruker Avance III-400 at 400 MHz ($^{1}$H NMR) and 100 MHz ($^{13}$C NMR) respectively by using deuterated solvents DMSO-$d_6$. The chemical shifts of $^{1}$H NMR (400 MHz) were taken relative to DMSO-$d_6$ as the internal reference (DMSO-$d_6$: δ = 2.50 ppm). The chemical shifts of $^{13}$C NMR (100 MHz) were measured using DMSO-$d_6$ as the internal standard (DMSO-$d_6$: δ = 39.52 ppm). Chemical shifts are denoted in parts per million (δ) and coupling constant (*J*) are reported in Hertz (Hz). The spin multiplicity are reported as singlet (s), doublet (d), triplet (t), quartet (q), broad singlet (bs), multiplet (m), doublet of doublet (dd), doublet of doublet of doublet (ddd), doublet of triplet (dt), and quartet of doublet (qd). LC-MS analysis was performed on a platform equipped with Agilent LC-MS 1260-6110 or Agilent LC-MS 1260-6120, using a Waters X Bridge C18: 50 mm x 4.6 mm x 3.5 um column. Flash column chromatography was conducted with silica gel (200-300 mesh, Qingdao Haiyang Chemical Co. Ltd., China). Analytical and preparative TLC analysis was performed on GF254 silica gel plates (Yantai Jiangyou Inc., China). Unless otherwise noted, reagents and all solvents are analytically pure grade and were obtained commercially from vendors such as Chron Chemical or Energy-Chemical. All compounds were determined to be >95% pure based on LC/MS and NMR.

##### 11.2. Chemical synthesis and Analytical Data of Compounds

All compounds were synthesized according to the reported method.[10] 4-Halo-6-substituted-quinoline (0.5 mmol, 1.0 equiv), N-(3-hydroxy-5-methoxyphenyl)acetamide (90.5 mg, 0.5 mmol, 1.0 equiv), and $K_2CO_3$ (138 mg, 1.0 mmol, 2.0 equiv) were added to a round-bottom flask equipped with a magnetic bar, then 3 mL of DMF was added as a solvent. The reaction vessel was evacuated and backfilled with $N_2$ three times and protected with a balloon of $N_2$. The reaction mixture was heated at 115 ℃ for at least 12 h with vigorous stirring. The solution was allowed to cool down to room temperature before being diluted with 20 mL ethyl acetate and washed with brine once. The organic phase was dried over anhydrous $Na_2SO_4$ and concentrated in vacuo. The residue was purified by silica gel flash chromatography to afford the corresponding product. **LG178, LG182, and LXY19,** which correspond to 10b,12a, and 12b, respectively, in our previous publication.[10]

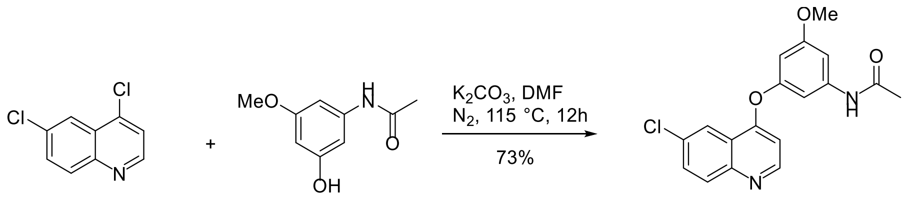


***N*-(3-((6-Chloroquinolin-4-yl)oxy)-5-methoxyphenyl)acetamide (SY142).** a white solid (124 mg, yield = 72.5%, purity = 98%) was prepared from 4,6-dichloroquinoline (118.2 mg, 0.6 mmol, 1.2 equiv) and N-(3-hydroxy-5-methoxyphenyl) acetamide (90.5 mg, 0.5 mmol, 1.0 equiv). TLC $R_f$ = 0.25 (PE/EA = 1/2). LCMS (m/z): calculated for $C_{18}H_{16}ClN_2O_3$ $[M+H]^+$ 343.08, found 343.50. **$^1$H NMR** (400 MHz, DMSO-$d_6$) δ 10.10 (s, 1H), 8.75 (d, *J* = 5.1 Hz, 1H), 8.26 (d, *J* = 2.4 Hz, 1H), 8.07 (d, *J* = 9.0 Hz, 1H), 7.85 (dd,

*J* = 9.1, 2.4 Hz, 1H), 7.15 (d, *J* = 2.4 Hz, 2H), 6.79 (d, *J* = 5.1 Hz, 1H), 6.62 (t, *J* = 2.2 Hz, 1H), 3.76 (s, 3H), 2.04 (s, 3H); **$^{13}$C NMR** (100 MHz, DMSO-$d_6$) δ 168.64, 160.92, 159.77, 154.76, 152.02, 147.63, 141.64, 131.06, 131.03, 130.77, 121.43, 120.30, 105.59, 103.29, 102.04, 101.18, 55.41, 39.49, 24.09.

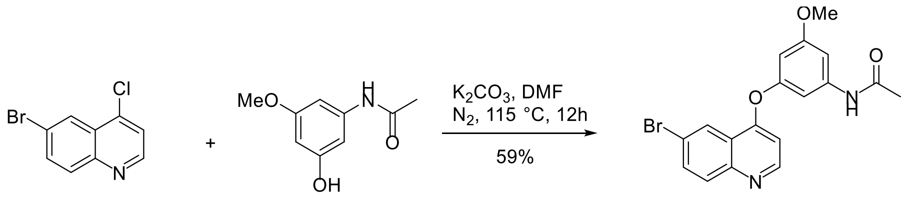


***N*-(3-((6-Bromoquinolin-4-yl)oxy)-5-methoxyphenyl)acetamide (LXY18).** a white solid (68 mg, yield = 58.5%, purity = 99%) was prepared from 6-bromo-4-chloroquinoline (72.75 mg, 0.3 mmol, 1.0 equiv) and N-(3-hydroxy-5-methoxyphenyl) acetamide (54.36 mg, 0.3 mmol, 1.0 equiv). TLC $R_f$ = 0.15 (PE/EA = 1/4). LCMS (m/z): calculated for $C_{18}H_{16}{}^{81}BrN_2O_3$ $[M+H]^+$ 389.02, found 388.90. **$^1$H NMR** (400 MHz, DMSO-$d_6$) δ 10.12 (s, 1H), 8.74 (d, *J* = 5.2 Hz, 1H), 8.41 (d, *J* = 2.2 Hz, 1H), 8.13 – 7.82 (m, 2H), 7.16 (d, *J* = 2.3 Hz, 2H), 6.77 (d, *J* = 5.2 Hz, 1H), 6.62 (t, *J* = 2.2 Hz, 1H), 3.76 (s, 3H), 2.04 (s, 3H); **$^{13}$C NMR** (100 MHz, DMSO-$d_6$) δ 168.64, 160.92, 159.64, 154.73, 152.12, 147.79, 141.65, 133.32, 131.12, 123.53, 121.92, 119.49, 105.55, 103.31, 102.06, 101.18, 55.41, 39.50, 24.09.

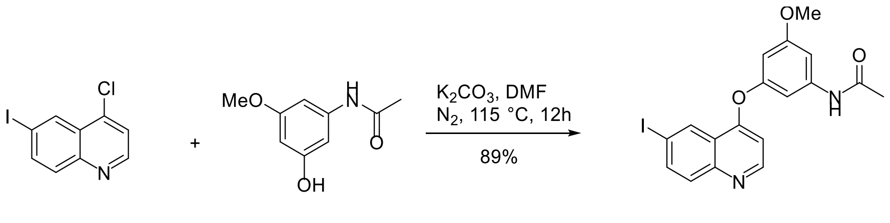

***N*-(3-((6-Iodoquinolin-4-yl)oxy)-5-methoxyphenyl)acetamide (LC08)**. a white solid (191.4 mg, yield = 88.2%, purity = 99%) was prepared from 4-chloro-6-iodoquinoline (173.7 mg, 0.6 mmol, 1.2 equiv) and N-(3-hydroxy-5-methoxyphenyl) acetamide (90.5 mg, 0.5 mmol, 1.0 equiv). TLC $R_f$ = 0.35 (PE/EA = 1/2). LCMS (m/z): calculated for $C_{18}H_{16}IN_2O_3$ $[M+H]^+$ 435.01, found 435.30. **$^1$H NMR** (400 MHz, DMSO-$d_6$) δ 10.10 (s, 1H), 8.74 (d, *J* = 5.2 Hz, 1H), 8.62 (d, *J* = 1.9 Hz, 1H), 8.09 (dd, *J* = 8.9, 2.0 Hz, 1H), 7.82 (d, *J* = 8.9 Hz, 1H), 7.15 (dt, *J* = 7.0, 2.0 Hz, 2H), 6.76 (d, *J* = 5.1 Hz, 1H), 6.62 (t, *J* = 2.2 Hz, 1H), 3.76 (s, 3H), 2.04 (s, 3H). **$^{13}$C NMR** (100 MHz, DMSO-$d_6$) δ 168.85, 161.14, 159.57, 154.97, 152.33, 148.27, 141.84, 138.82, 131.02, 130.10, 122.61, 105.63, 103.51, 102.27,

#### 11.3. Spectral Data of Compounds (figures)

$^{1}$H NMR spectrum (400 MHz, DMSO-$d_6$) of compound **SY142**

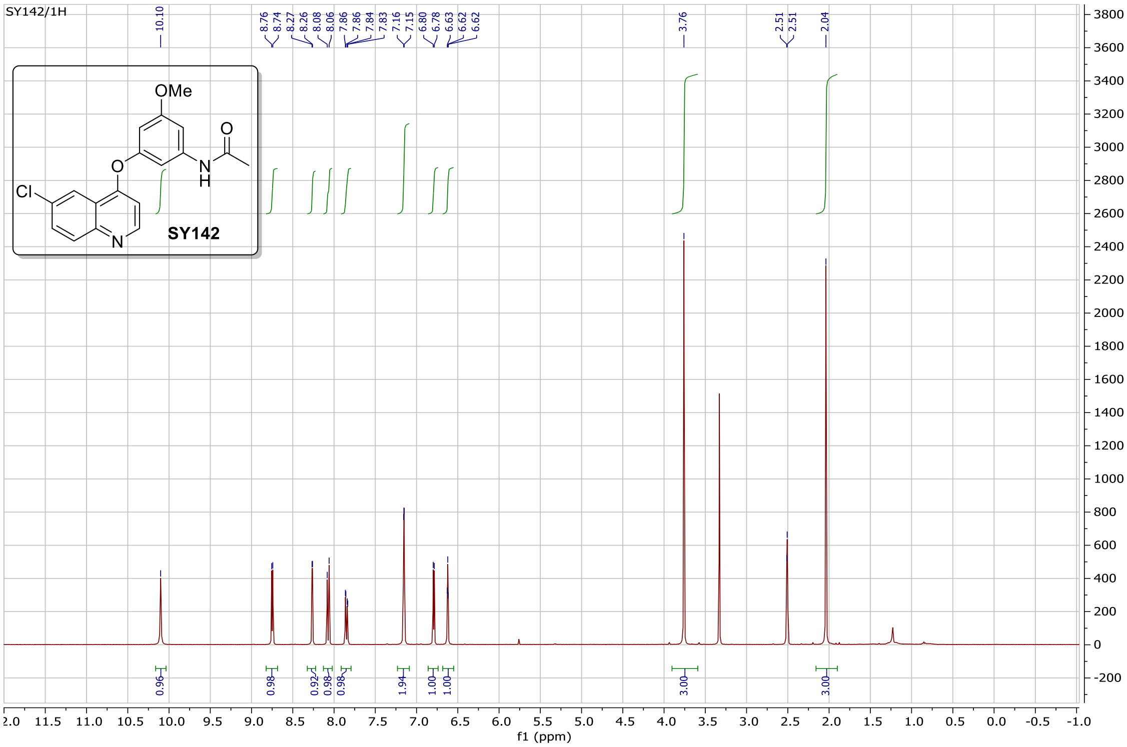


$^{13}$C NMR spectrum (100 MHz, DMSO-$d_6$) of compound **SY142**

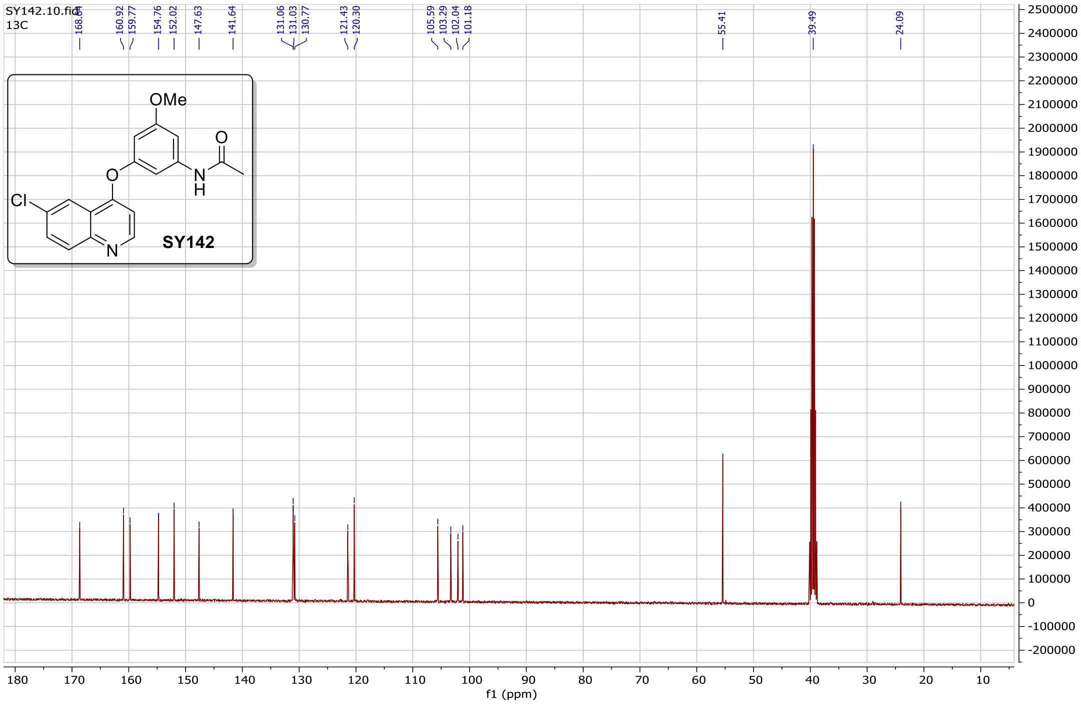

#### $^{1}$H NMR spectrum (400 MHz, DMSO-$d_6$) of compound **LXY18**

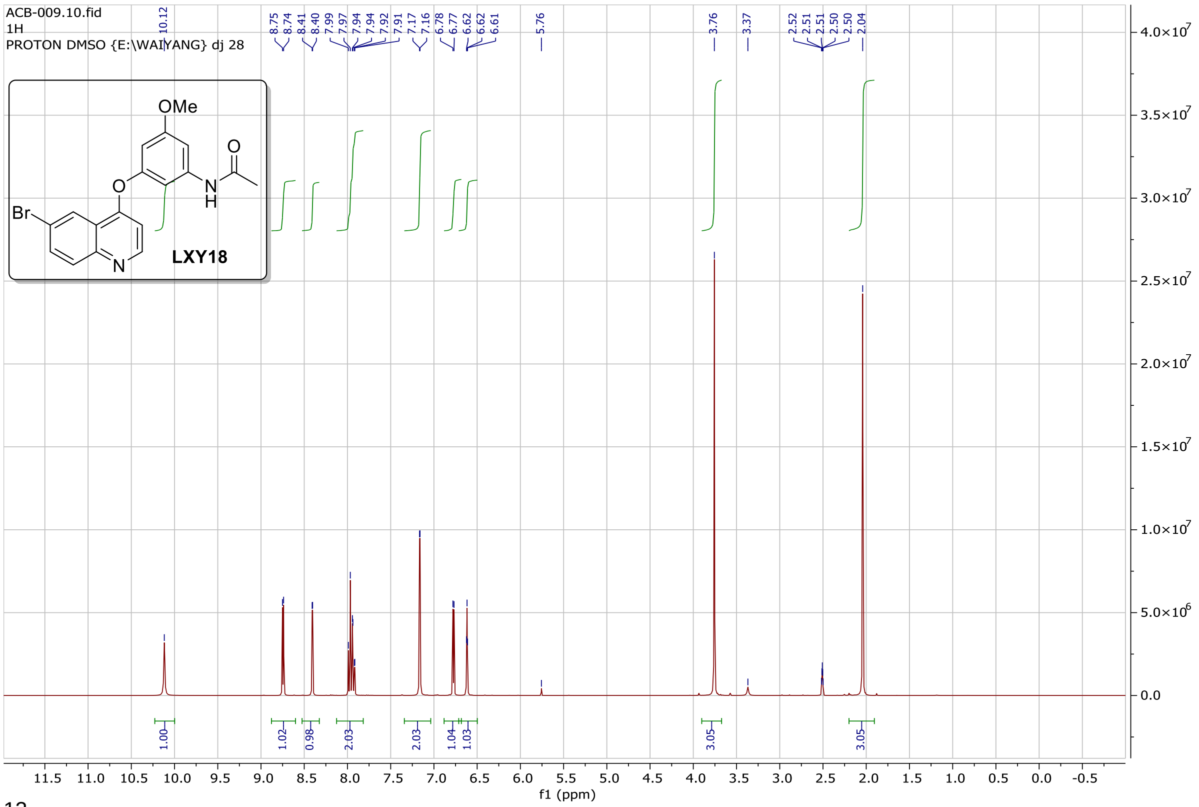

#### $^{13}$C NMR spectrum (100 MHz, DMSO-$d_6$) of compound **LXY18**

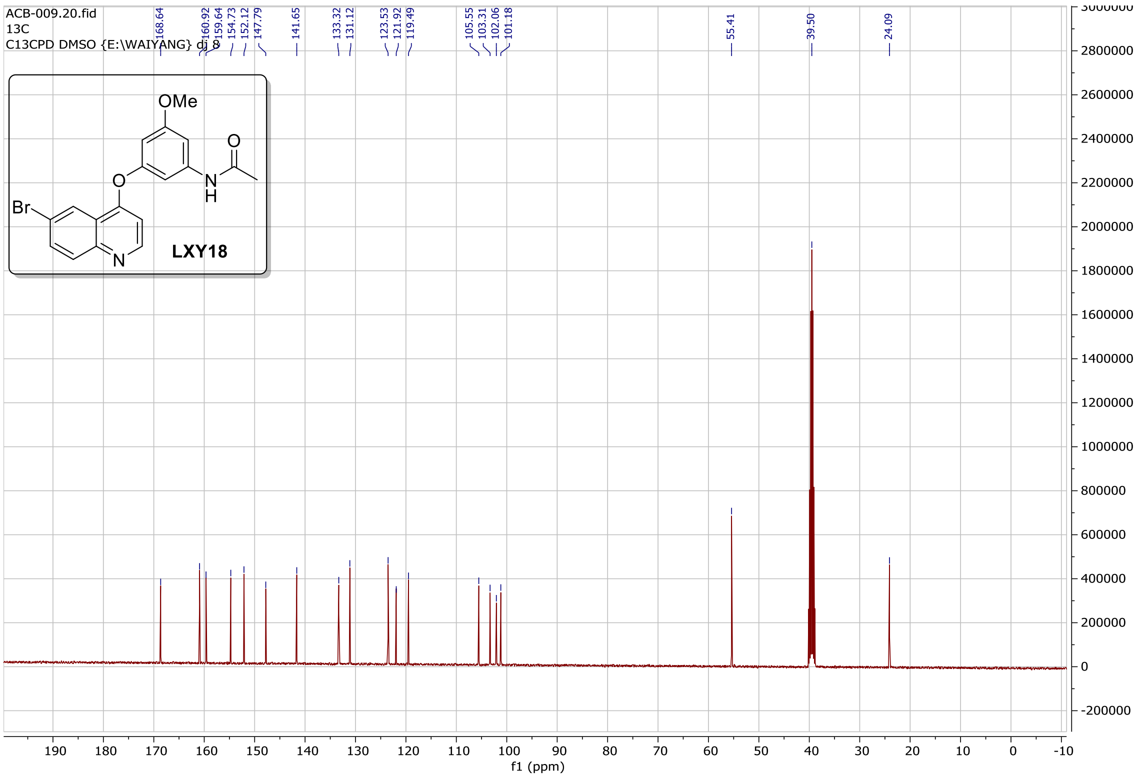

#### $^{1}$H NMR spectrum (400 MHz, DMSO-$d_6$) of compound **LC08**

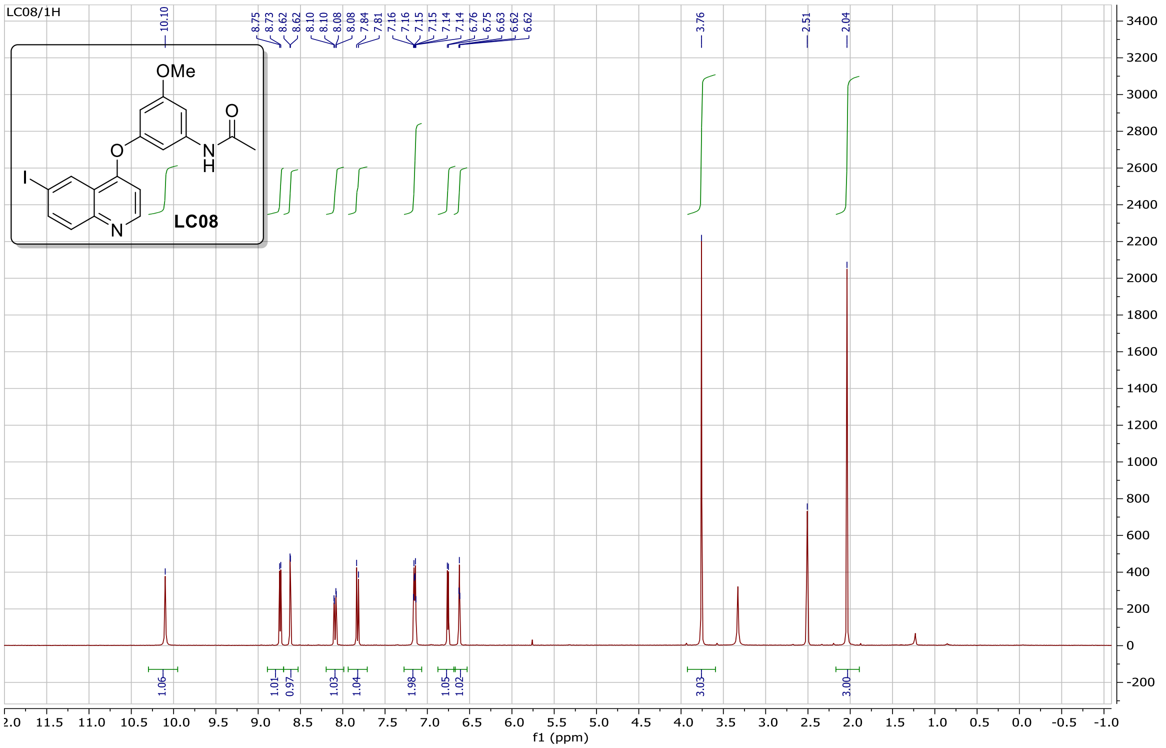


#### $^{13}$C NMR spectrum (100 MHz, DMSO-$d_6$) of compound **LC08**

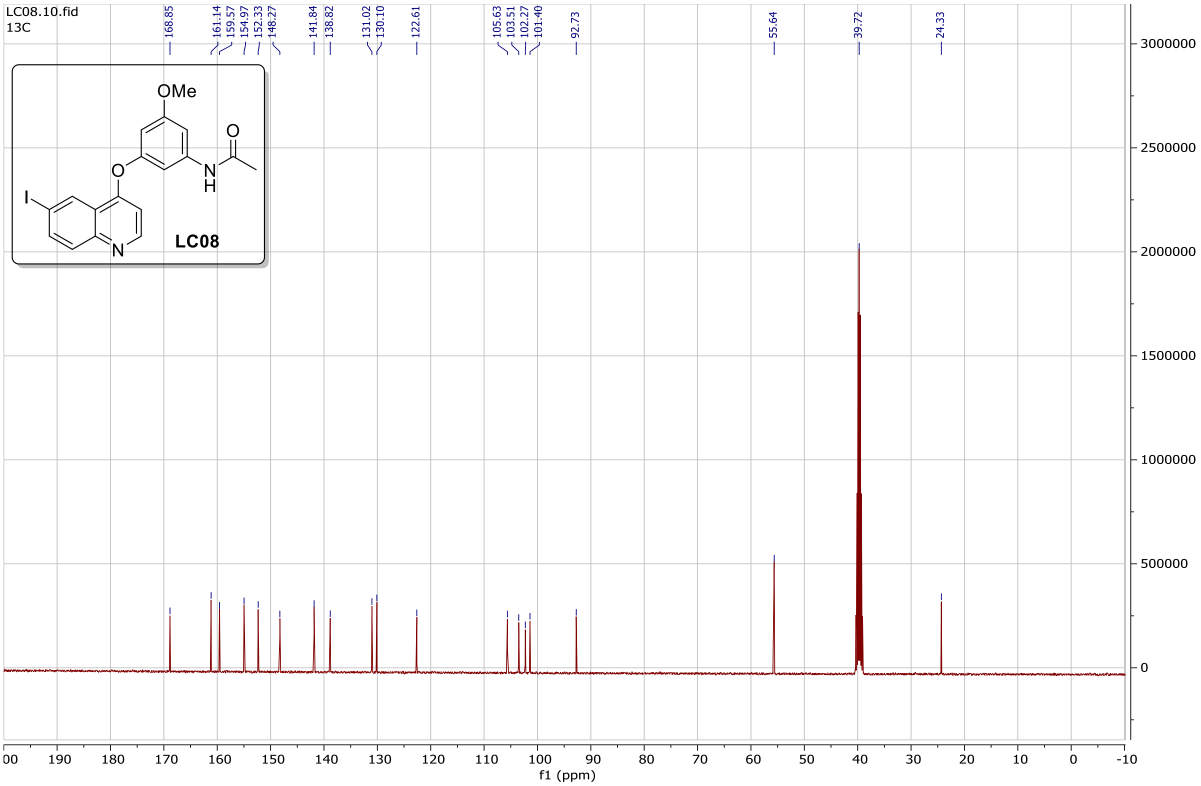

(7) Daina, A.; Michielin, O.; Zoete, V. SwissADME: A Free Web Tool to Evaluate Pharmacokinetics, Drug-Likeness and Medicinal Chemistry Friendliness of Small

Molecules. *Sci Reports 2017 71* **2017**, *7* (1), 1–13.

(8) Eriksson, M. A. L.; Gabrielsson, J.; Nilsson, L. B. Studies of Drug Binding to Plasma Proteins Using a Variant of Equilibrium Dialysis. *J Pharm Biomed Anal* **2005**, *38* (3), 381–389.

(9) Lv, G.; Shi, Q.; Zhang, T.; Li, J.; Kalashova, J.; Long, Y.; Sun, Y.; Li, C.; Choudhry, N.; Li, H.; Yang, C.; Zhou, X.; Reddy, M. C.; Anantoju, K. K.; Jupelli, R.; Zhang, S.; Zhang, J.; Allen, T.; Liu, H.; Nimishetti, N.; Yang, D. 2-Phenoxy-3, 4′-Bipyridine Derivatives Inhibit AURKB-Dependent Mitotic Processes by Disrupting Its Localization. *Eur J Med Chem* **2022**, 114904.

(10) Lv, G.; Shi, Q.; Zhang, T.; Li, J.; Long, Y.; Zhang, W.; Choudhry, N.; Yang, K.; Li, H.; Kalashova, J.; Yang, C.; Zhou, X.; Chenna Reddy, M.; Kumar Anantoju, K.; Zhang, S.; Zhang, J.; Allen, T. D.; Liu, H.; Nimishetti, N.; Yang, D. Integrating a Phenotypic Screening with a Structural Simplification Strategy to Identify 4-Phenoxy-Quinoline Derivatives to Potently Disrupt the Mitotic Localization of Aurora Kinase B. *Bioorg Med Chem* **2023**, 117173.
